# Mathematical Relationships between Spinal Motoneuron Properties

**DOI:** 10.1101/2021.08.05.455188

**Authors:** Arnault Caillet, Andrew T.M. Phillips, Dario Farina, Luca Modenese

**Affiliations:** Department of Civil and Environmental Engineering, Imperial College London, SW7 2AZ, UK; Department of Bioengineering, Imperial College London, SW7 2AZ, UK; Graduate School of Biomedical Engineering, University of New South Wales, Sydney, Australia

**Keywords:** motoneuron, motoneuron size, motoneuron membrane properties, motor unit, Henneman’s size principle

## Abstract

Our understanding of the behaviour of spinal alpha-motoneurons (MNs) in mammals partly relies on our knowledge of the relationships between MN membrane properties, such as MN size, resistance, rheobase, capacitance, time constant, axonal conduction velocity and afterhyperpolarization period.

We reprocessed the data from 40 experimental studies in adult cat, rat and mouse MN preparations, to empirically derive a set of quantitative mathematical relationships between these MN electrophysiological and anatomical properties. This validated mathematical framework, which supports past findings that the MN membrane properties are all related to each other and clarifies the nature of their associations, is besides consistent with the Henneman’s size principle and Rall’s cable theory.

The derived mathematical relationships provide a convenient tool for neuroscientists and experimenters to complete experimental datasets, to explore relationships between pairs of MN properties never concurrently observed in previous experiments, or to investigate inter-mammalian-species variations in MN membrane properties. Using this mathematical framework, modelers can build profiles of inter-consistent MN-specific properties to scale pools of MN models, with consequences on the accuracy and the interpretability of the simulations.

## Introduction

Assessing the morphological and electrophysiological properties of individual spinal alpha-motoneurons (MNs) is crucial for understanding the recruitment and discharge behaviour of MNs and for exploring the neuro-mechanical interplay that controls voluntary motion in mammals. As measured in individual spinal MNs in experimental studies and reported in review papers (Henneman, 1981; Burke, 1981; Binder et al., 1996; Powers & Binder, 2001; Kernell, 2006; Heckman & Enoka, 2012), significant correlations exist in mammalian MN pools between the MN morphological and/or electrophysiological properties reported in Table 1. For example, the soma size (*D_soma_*) and the current threshold for spike initiation (*I_th_*) of a MN are both positively correlated to the axonal conduction velocity (*ACV*) and afterhyperpolarization period (*AHP*). Also, *I_th_* decreases with increasing input resistance (*R*) and *AHP, R* is negatively correlated to *ACV* and *D_soma_*, and *ACV* varies inversely with *AHP*. Besides supporting and sequentially leading to extensions of the Henneman’s size principle (Henneman, 1957; Wuerker et al., 1965; Henneman et al., 1965a; Henneman et al., 1965b; Henneman et al., 1974; Henneman, 1981; Henneman, 1985), these empirical associations find strong consistency with Rall’s theoretical approach of representing the soma and the dendritic trees of MNs as an equivalent cylinder (Rall, 1977; Rall et al., 1992; Powers & Binder, 2001). In such case, MNs behave like resistive-capacitive (RC) circuits, as successfully simulated with computational RC models (Izhikevich, 2004; Dong et al., 2011; Negro et al., 2016; Teeter et al., 2018), that rely on further property associations, such as the MN membrane time constant *τ* precisely equalling the product of the MN membrane specific capacitance *C_m_* and resistivity *R_m_* (*τ* = *R_m_C_m_*).

**Table 1.** The motoneuron (MN) and muscle unit (mU) properties investigated in this study with their notations and SI base units. *S_MN_* is the size of the MN. As reproduced in Table 2, the MN size *S_MN_* is adequately described by measures of the MN surface area *S_neuron_* and the soma diameter *D_soma_*. *R* and *R_m_* define the MN-specific electrical resistance properties of the MN and set the value of the MN-specific current threshold *I_th_* (Binder et al., 1996; Powers & Binder, 2001; Heckman & Enoka, 2012). C and *C_m_* (constant among the MN pool) define the capacitance properties of the MN and contribute to the definition of the MN membrane time constant τ (Gustafsson & Pinter, 1984a; Zengel et al., 1985). Δ*V_th_* is the amplitude of the membrane voltage depolarization threshold relative to resting state required to elicit an action potential. *I_th_* is the corresponding electrical current causing a membrane depolarization of Δ*V_th_*. AHP is defined in most studies as the duration between the action potential onset and the time at which the MN membrane potential meets the resting state after being hyperpolarized. ACV is the axonal conduction velocity of the elicited action potentials on the MN membrane. *S_mU_* is the size of the mU. As indicated in Table 2, the mU size *S_mU_* is adequately described by measures of (1) the sum of the cross-sectional areas (CSAs) of the fibres composing the mU *CSA^tot^*, (2) the mean fibre CSA *CSA^mean^*, (3) the innervation ratio IR, i.e. the number of innervated fibres constituting the mU, and (4) the mU tetanic force *F^tet^*. *F^tw^* is the MU twitch force.

|  | Properties | Notation | Unit |
| --- | --- | --- | --- |
| **Motoneuron (MN) properties** | Size: | $S_{MN}$ | |
| | Neuron surface area | $S_{neuron}$ | $[m^2]$ |
| | Soma diameter | $D_{soma}$ | $[m]$ |
| | Resistance | $R$ | $[\Omega]$ |
| | Specific resistance per unit area | $R_m$ | $[\Omega \cdot m^2]$ |
| | Capacitance | $C$ | $[F]$ |
| | Specific capacitance per unit area | $C_m$ | $[F \cdot m^{-2}]$ |
| | Time constant | $\tau$ | $[s]$ |
| | Rheobase (current recruitment threshold) | $I_{th}$ | $[A]$ |
| | Voltage threshold | $\Delta V_{th}$ | $[V]$ |
| | Afterhyperpolarization period | $AHP$ | $[s]$ |
| | Axonal conduction velocity | $ACV$ | $[m \cdot s^{-1}]$ |
| **Muscle unit (mU) properties** | Size: | $S_{mU}$ | |
| | Total fibre cross-sectional area (CSA) | $CSA^{tot}$ | $[m^2]$ |
| | Mean fibre CSA | $CSA^{mean}$ | $[m^2]$ |
| | Innervation ratio | $IR$ | $[ ]$ |
| | Tetanic force | $F^{tet}$ | $[N]$ |
| | Twitch force | $F^{tw}$ | $[N]$ |

Yet, because the empirical correlations between MN properties were obtained from scattered data from individual experimental studies, the quantitative mathematical associations between the MN properties reported in Table 1, beyond their aforementioned global relative variations, remain unclear. For example, the negative correlation between *R* and *ACV* was described in the literature with linear (Fleshman et al., 1981), exponential (Burke, 1968) or power relationships (Kernell, 1966; Kernell & Zwaagstra, 1981; Ulfhake & Kellerth, 1984; Gustafsson & Pinter, 1984a), while Kernell & Zwaagstra (1980) reported for this association a slope twice greater than Gustafsson & Pinter (1984a) in the double-logarithmic space. This makes it difficult to reconcile the conclusions of multiple empirical studies that investigated different property associations. For example, Burke et al. (1968) and Zwaagstra et al. (1980) respectively proposed exponential *R* – *ACV* and linear *ACV* – *AHP* associations, suggesting a hybrid exponential relationship between *R* and *AHP*, that is different from the power *R – AHP* relationship directly reported by Ulfhake & Kellerth (1984). These divergences between studies are a major limitation for our understanding of the associations and the distribution of MN properties in a MN pool, which cannot be directly investigated experimentally either, as measuring multiple MN morphometric properties *in vitro* and electrophysiological properties *in vivo* for a large sample of MNs is challenging.

Here, we reprocessed and merged the published data from 19 available experimental studies in adult cat preparations to derive and validate a unique set of mathematical power relationships between all pairs of the MN morphometric and electrophysiological cat properties listed in Table 1. The significance of these quantitative relationships, which are consistent with the conclusions from Rall’s cable theory, supports the notion that the properties reported in Table 1 are all associated to each other. The uniqueness of these mathematical relationships tackles the aforementioned inter-study variability in describing the data and clarifies our understanding of the quantitative association between spinal MN properties, including for pairs of properties never simultaneously measured in experiments. These relationships also provide a convenient mathematical framework for modelers for the derivation of appropriate and consistent MN profiles of MN-specific morphometric and electrophysiological properties for the realistic scaling of pools of computational models of MNs, improving the interpretability of model predictions. Using the relationships, experimenters can readily complete their datasets by deriving MN-specific values, that are representative of the literature, for the MN properties that were not experimentally measured.

After deriving the mathematical framework from cat data, we then demonstrate that the normalized cat relationships apply to other mammals by validating them with data from 9 adult rat and mouse electrophysiological studies *in vivo*. This approach helps better understanding the inter-mammalian-species variations in MN properties. Finally, using additional correlations obtained between some MN and muscle unit (mU) properties from 14 mammal studies, we discuss the empirical relationships obtained between MN properties in the context of the Henneman’s size principle.

## Methods

### Selected studies reporting processable adult mammal data

#### Identification of the selected studies

To optimize inter-study consistency, we only selected studies that concurrently measured at least two of the morphometric and/or electrophysiological properties reported in Table 1 in individual spinal alpha-motoneurons of healthy adult cats, rats and mice. To build an extensive set of relevant studies so that the mathematical relationships derived in this study describe a maximum of the data published in the literature, the output of the systematic analysis provided in Highlander et al. (2020) was screened and used for a further search by reference lists. Among the retrieved studies, larval and postnatal specimens were disregarded because of the pronounced age-related variance (Highlander et al., 2020) in the mean values of the electrophysiological properties listed in Table 1, which corroborates the non-extrapolability of neonatal neuronal circuitry to older ages (Song et al., 2006; Carp et al., 2008; Nakanishi & Whelan, 2010; Mitra & Brownstone, 2012). Due to the nonlinear inter-species scalability of the spinal alpha-MN electrophysiological and morphometric properties (Manuel et al., 2019) and a pronounced variance in the inter-species mean values (Highlander et al., 2020), non-mammalian species were also disregarded, while the retained data from cats, rats and mice were processed separately. Highlander et al. (2020) also report an ‘unexplained’ variance in the mean electrophysiological property values reported between studies investigating specimens of the same species, sex and age, which may be explained by differences between *in vivo* and *in vitro* protocols (Carp et al., 2008). For this reason, only studies measuring the electrophysiological properties *in vivo* were considered, while most *in vitro* studies had already been disregarded at this stage as dominantly performed on neonatal specimens due to experimental constraints (Manuel et al., 2009; Mitra & Brownstone, 2012). As all but two of the remaining selected studies focused on lumbosacral MNs, the final set of considered publications was constrained to MNs innervating hindlimb muscles to improve inter-study consistency. From a preliminary screening of the final set of selected studies, it was finally found that relatively more cat studies (19) were obtained than rat (9) and mouse (4) studies, while the cat studies also investigated more pairs of morphometric and electrophysiological properties. The mathematical relationships sought in this paper between MN properties were thus derived from the cat data reported in the selected studies, and then validated for extrapolation to rat and mouse data.

#### Selected studies providing cat data – experimental approaches

The 19 selected studies focusing on cats that were included in this work were published between 1966 and 2001. They applied similar experimental protocols to measure the morphometric and electrophysiological properties reported in Table 1. All the selected studies performing morphometric measurements injected *in vitro* the recorded MNs intracellularly with horseradish peroxidase (HRP) tracer by applying a continuous train of anodal current pulses. The spinal cord was eventually sliced frozen or at room temperature with a microtome or a vibratome and the MN morphometry was investigated. All morphometric measurements were manually performed, and the MN compartments (soma, dendrites, axon) approximated by simple geometrical shapes, yielding some experimental limitations discussed later in the Methods and Supplementary Material sections.

To measure the electrophysiological properties, animals were anesthetized, immobilized, and kept under artificial breathing. Hindlimb muscles of interest were dissected free, and their nerves were mounted onto stimulating electrodes, while maintaining body temperature between 35°C and 38°C. After a laminectomy was performed over the lumbosacral region of the spinal cord, the MNs were identified by antidromic invasion following electrical stimulation of the corresponding muscle nerves. All selected cat studies reported the use of electrodes having stable resistances and being able to pass currents up to 10 nA without evident polarization or rectification. Some aspects of the experimental protocol however diverged between the selected studies, such as the age and the sex of the group of adult cats, the size population of cats and recorded MNs, the muscles innervated by the recorded MNs, the means of anaesthesia, the level of oxygen, and whether the spinal dorsal roots were severed (Kernell, 1966; Burke, 1968; Barrett & Crill, 1974; Fleshman et al., 1981; Gustafsson & Pinter, 1984a; Gustafsson & Pinter, 1984b; Zengel et al., 1985; Foehring et al., 1987), the complete surrounding hindlimbs were denervated (Burke, 1968; Kernell & Zwaagstra, 1981) or the spinal cord was transected (Burke, 1968; Krawitz et al., 2001).

In all the selected cat studies, *ACV* was calculated as the ratio of the conduction distance and the antidromic spike latency between the stimulation site and the ventral root entry. The studies however did not report how the nerve length was estimated. *AHP* was measured using brief suprathreshold antidromic stimulations, as the time interval between the spike onset to when the voltage deflection of the hyperpolarizing phase returned to the prespike membrane potential. In all studies, *I_th_* was obtained as the minimal intensity of a 50-100ms-long depolarizing rectangular current pulse required to produce intermittent discharges. *I_th_* was obtained with a trial-and-error approach, slowly increasing the intensity of the pulse. The selected studies did not report any relevant source of inaccuracy in measuring *ACV, AHP* and *I_th_*. The studies measured *R* using the spike height method (Frank & Fuortes, 1956). In brief, small (1-5nA) depolarizing and/or hyperpolarizing current pulses (*I_in_*) of 15-100ms duration were injected through a Wheatstone bridge circuit in the intracellular microelectrode amplifier, and the subsequent change in the amplitude of the membrane voltage potential (*V_m_*) was reported in steady-state *V_m_* – *I_in_* plots. In all studies, *R* was obtained by calculating the slope of the linear part of the *V_m_* – *I_in_* plots near resting membrane potential, by analogy with Ohm’s law. It is worth noting for inter-study variability that depolarizing pulses return slightly higher *R* values than hyperpolarizing currents (Sasaki, 1991). To measure the membrane time constant *τ*, all the selected studies analysed the transient voltage responses (*V_m_*) to weak (1-12nA) brief (0.5ms) or long (15-100ms) hyperpolarizing current pulses. Both brief- and long-pulse approaches were reported to provide similar results for the estimation of *τ* (Ulfhake & Kellerth, 1984). Considering MNs as equivalent isopotential cables, the membrane time constant *τ* was identified in all studies as the longest time constant when modelling the measured voltage transient response to a current pulse as the sum of weighted time-dependent exponential terms (Rall, 1969; Rall, 1977; Powers & Binder, 2001). Semi-logarithmic plots of the time-history of the voltage transient *V_m_* or of its time-derivative 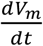 were drawn, and a straight line was fit by eye to the linear tail of the resulting plot (Fleshman et al., 1988), the slope of which was *τ*. A graphical ‘peeling’ process (Rall, 1969) was undertaken to recover the first equalizing time constant *τ*_1_, required to estimate the electronic length (*L*) of Rall’s equivalent cylinder representation of the MN and the MN membrane capacitance with Rall’s equations (Rall, 1969; Rall, 1977; Gustafsson & Pinter, 1984b; Powers & Binder, 2001):

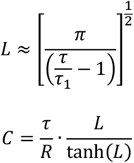

These electrophysiological measurements were reported to be subject to three main experimental sources of inaccuracy. First, a variable membrane ‘leak’ conductance, that can be estimated from the ratio of two parameters obtained from the ‘peeling’ process (Gustafsson & Pinter, 1984b), arises from the imperfect seal around the recording micropipette (Ulfhake & Kellerth, 1984; Gustafsson & Pinter, 1984b; Pinter & Vanden Noven, 1989). As reviewed in Powers & Binder (2001) and in Kernell (2006), this ‘leak’ can affect the measurements of all the properties, notably underestimating the values of *R* and overestimating those of *τ* and *C*. However, this ‘leak’ is probably not of major significance in the cells of large spike amplitudes (Gustafsson & Pinter, 1984b), on which most of the selected studies focused. Importantly, the membrane ‘leak’ is also not expected to affect the relations between the parameters (Gustafsson & Pinter, 1984b).

Secondly, some nonlinearities (Ito & Oshima, 1965; Burke & Ten Bruggencate, 1971; Ulfhake & Kellerth, 1984) in the membrane voltage response to input current steps arise near threshold because of the contribution of voltage-activated membrane conductances to the measured voltage decay (Fleshman et al., 1988). This contradicts the initial assumption of the MN membrane remaining passive to input current steps (Rall, 1969; Burke & Ten Bruggencate, 1971). Because of this issue, the ends of the *V_m_* – *I_in_* plot are curvilinear and can affect the estimation of *R*, while the transient voltage response to an input current pulse decays faster than exponentially (it undershoots) at the termination of the current injection, which makes it impossible to plot the entire course of ln(*V*(*t*)) and challenges the graphical procedure taken to estimate *τ* (Fleshman et al., 1988). Therefore, some of the selected studies discarded all the MNs that displayed obvious large nonlinearity in the semi-logarithmic V-t or 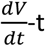 plots (Burke & Ten Bruggencate, 1971). The membrane nonlinearities can be corrected by adding a constant voltage to the entire trace (Fleshman et al., 1988) or by considering the three time-constant model of Ito & Oshima (1965) to approximate and subtract to the voltage signal the membrane potential change produced by the current steps. However, none of the selected studies performed such corrections, potentially yielding a systematic underestimation of the values of *R* and *τ* (Gustafsson & Pinter, 1984b; Zengel et al., 1985; Powers & Binder, 2001). Yet, this systematic error for *R*, which should not contribute to inter-study variability, is expected to remain low because the selected studies used input current pulses of low strength (Ulfhake & Kellerth, 1984) and measured *R* from the linear part of the I-V plots near the resting membrane potential where no membrane nonlinearity is expected to occur. This systematic error may also not affect the distribution of recorded *R* values, as displayed in Zengel et al. (1985). Zengel et al. (1985) besides corrected for the membrane nonlinearities with the three time-constant model approach of Ito & Oshima (1965) when measuring *τ*, and returned values of *τ* around 40% higher than the other selected studies. However, this systematic discrepancy is expected to disappear with the normalization of the datasets described in the following, as the reported normalized distributions of *τ* are very similar between the selected studies (see density histogram for the {*τ*; *R*} dataset in Figure FSM1 in Supplementary Material). Thirdly, the highest source of inaccuracy and inter-study variability arises from the subjective fit ‘by eye’ of a straight line to the transient voltage when estimating *τ* (Pinter & Vanden Noven, 1989).

Overall, all the selected studies used similar experimental approaches to measure *ACV, AHP, I_th_, R, C* and *τ*, and little sources of inaccuracy and inter-study variability were identified.

### Relationships between MN properties

For convenience, in the following, the notation {*A*; *B*} refers to the pair of morphometric or electrophysiological MN properties *A* and *B*, with *A, B* ∈ {*S_MN_*; *ACV*; *AHP*; *R*; *I_th_*; *C*; *τ*}, defined in Table 1. The selected studies generally provided clouds of data points for pairs {*A*; *B*} of concurrently measured MN properties through scatter graphs. These plots were manually digitized using the online tool WebPlotDigitizer (Ankit, 2020). When a study did not provide such processable data – most reported the mean ±*SD* property values of the cohort of measured MNs – the corresponding author was contacted to obtain the raw data of the measured MN properties, following approval of data sharing. Upon reception of data, datasets of all possible {*A*; *B*} pairs were created and included into the study.

#### Normalized space and choice of regression type

The sets of data retrieved from different cat studies for each property pair {*A*; *B*} were merged into a ‘global’ dataset dedicated to that property pair. The {*A*; *B*} data was also normalized for each study and transformed as a percentage of the maximum property value measured in the same study and normalized ‘global’ datasets were similarly created. A least squares linear regression analysis was performed for the ln(*A*) – ln(*B*) transformation of each global dataset yielding relationships of the type ln(*A*) = *a* · ln(*B*) + *k*, which were then converted into power relationships of the type *A* = *k* · *B^a^* (eq.1). The adequacy of these global power trendlines and the statistical significance of the correlations were assessed with the coefficient of determination *r*^2^ (squared value of the Pearson’s correlation coefficient) and a threshold of 0.01 on the *p*-value of the regression analysis, respectively. For each {*A*; *B*} pair, the normalized global datasets systematically returned both a higher *r*^2^ and a lower *p*-value than the datasets of absolute {*A*; *B*} values, in agreement with the ‘unexplained’ inter-experimental variance reported in (Highlander et al., 2020). It was therefore decided to investigate the {*A*; *B*} pairs of MN properties in the normalized space using the normalized global datasets to improve the cross-study analysis. For this preliminary analysis and the rest of the study, power regressions (eq.1) were preferred to linear (*A* = *k* + *a* · *B*) or exponential (*A* = *k* · *e^a·B^*) regressions to maintain consistency with the mathematical structure of the equations from Rall’s cable theory (Rall, 1957; Rall, 1959; Rall, 1960; Powers & Binder, 2001) and of other well-defined relationships, such as 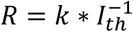. Also, a least squares linear regression analysis was preliminary performed for each normalized experimental dataset and for its ln(*A*) – ln(*B*) and ln(*A*) – *B* transformations, yielding linear, power, and exponential fits to the data. The power regressions overall returned *r*^2^ > 0.5 for more experimental datasets than the linear and exponential fits (see Table TSM1 in Supplementary Material). To avoid bias, power regressions were the only type of fit used in this study, despite a few datasets being more accurately fitted by linear or exponential regressions (the difference in the quality of the fit was very small in all cases). Other regression types, such as polynomial fits, were not justified by previous findings in the literature.

#### Global datasets and data variability

The data variability between the studies constituting the same global dataset {*A*; *B*} was assessed by analysing the normalized distributions of the properties *A* and *B* with four metrics, that were the range of the measured data, the mean of the distribution, the coefficient of variation (CoV) calculated as the standard deviation (sd) divided by the mean of the distribution, and the ratio 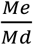 of the median and the mean of the distribution. The inter-study data variability in the {*A*; *B*} global dataset was assessed by computing the *mean_g_* ± *sd_g_* across studies of each of the four metrics. Low variability between the data distributions from different studies was concluded if the normalized distributions (in percentage maximum value) returned similar values for the range, mean, CoV and 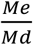 metrics, i.e. when *sd_g_* < 10% and 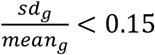 were obtained for all four metrics. In such case, the data distributions would respectively span over similar bandwidth length of the MU pool, be centred around similar mean values, display similar data dispersion and similar skewness. The relative size of the experimental datasets reported by the selected studies constituting the same global datasets was also compared to assess their relative impact on the regression curves fitted to the global datasets. Then, the inter-study variability in associating the distributions of properties *A* and *B* was assessed by computing the 95% confidence interval of the linear model fitted to the global dataset {*A*; *B*} in the log-log space, that yields the power regression *A* = *k* · *B^a^*. Low inter-study variability was considered if the value of *a* varied less than 0.4 in the confidence interval.

The variability of the data distribution of a property *A* between multiple global datasets {*A*; *B*} was then assessed by computing the same four metrics as previously (range, mean, CoV, 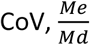) for each global dataset, and then computing the *mean_G_* ± *sd_G_* of each of the four metrics across the global datasets {*A*; *B*}. Low variability between the global datasets in the distribution of property *A* was considered if *sd_G_* < 10% and 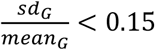 were obtained for all four metrics. If the inter-study and inter-global datasets variability was low, the global datasets were created and processed to derive the mathematical relationships between MN properties according to the procedure described below.

#### Size-dependent normalized relationships

From inspection of the considered cat studies, most of the investigated MN property pairs comprised either a direct measurement of MN size, noted as *S_MN_* in this study, or another variable well-admitted to be strongly associated to size, such as *ACV*, *AHP* or *R*. Accordingly, and consistently with Henneman’s size principle, we identified *S_MN_* as the reference MN property with respect to which relationships with the electrophysiological MN properties in Table 1 were investigated. To integrate the data from all global normalized datasets into the final relationships, the MN properties in Table 1 were processed in a step-by-step manner in the order *ACV, AHP, R, I_th_, C, τ*, as reproduced in Figure 1 for two arbitrary properties, to seek a ‘final’ power relationship of the type eq.1 between each of them and *S_MN_*. For each electrophysiological property *A* and each {*A*; *B*} dataset, we considered two cases. If *B* = *S_MN_*, the global dataset was not further processed as the electrophysiological property *A* was already related to MN size *S_MN_* from experimental measurements. If *B* ≠ *S_MN_*, the global {*A*; *B*} dataset was transformed into a new {*A*; *S_MN_*} dataset by converting the discrete values of *B* with the trendline regression *S_MN_* = *k_d_* · *B^d^*, which was obtained at a previous step of the data processing. With this dual approach, as many MN size-dependent {*A*; *S_MN_*} intermediary datasets as available {*A*; *B*} global datasets were obtained for each property *A*. These size-dependent datasets were merged into a ‘final’ {*A*; *S_MN_*} dataset to which a least squares linear regression analysis was performed for the ln(*A*) – ln(*S_MN_*) transformation, yielding relationships of the type ln(*A*) = *c* · ln(*S_MN_*) + *k_c_*, which were converted into the power relationships:

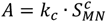

**Figure 1.**
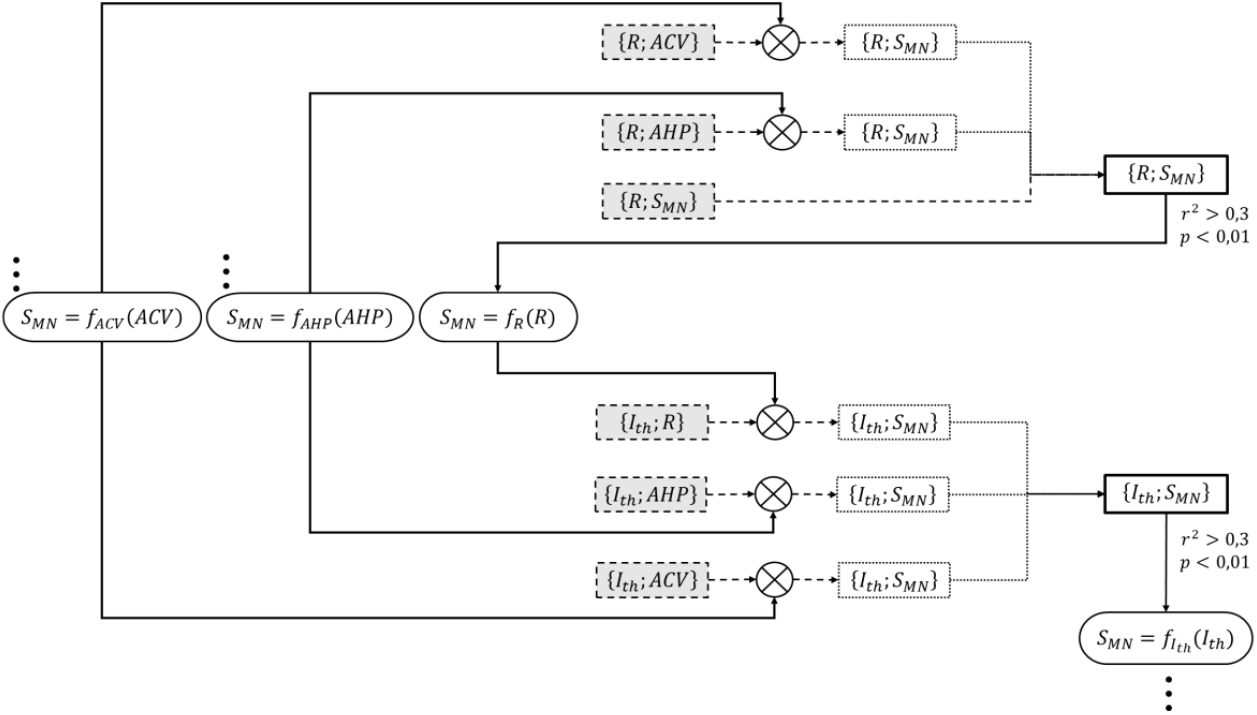
Detailed example for the process adopted to successively create the two final {*R; S_MN_*} and {*I_th_*; *S_MN_*} datasets. (right-side thick-solid contour rectangular boxes). These final datasets were obtained from respectively three and three normalized global datasets of experimental data obtained from the literature (dashed-contour grey-filled boxes) {{*R*; *ACV*}, {*R*; *AHP*}, {*R*; *S_MN_*}} and {{*I_th_*; *R*}, {*I_th_*; *AHP*}, {*I_th_*; *ACV*}}. The {*R*; *ACV*} and {*R*; *AHP*} datasets were first transformed (⊗ symbol) into two intermediary {*R*; *S_MN_*} datasets (dotted-contour boxes) by converting the *ACV* and *AHP* values to equivalent *S_MN_* values with two ‘inverse’ *S_MN_* = *f_ACV_*(*ACV*) and *S_MN_* = *f_AHP_*(*AHP*) power relationships (oval boxes with triple dots), that had been previously obtained from two unshown steps that had yielded the final {*ACV*; *S_MN_*} and {*AHP*; *S_MN_*} datasets. The two intermediary {*R*; *S_MN_*} datasets were merged with the remaining global {*R*; *S_MN_*} dataset to yield the final {*R*; *S_MN_*} dataset, to which a power relationship of the form 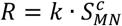 was fitted. If *r*^2^ > 0.3 and *p* < 0.01, an ‘inverse’ power relationship *S_MN_* = *f_R_*(*R*) (oval box) was further fitted to this final dataset. In a similar approach, the three normalized global datasets {*I_th_*; *R*}, {*I_th_*; *AHP*} and {*I_th_*; *ACV*} were transformed with the three ‘inverse’ relationships into intermediary {*I_th_*;*S_MN_*} datasets, which were merged to yield the final {*I_th_*;*S_MN_*} dataset. An ‘inverse’ *S_MN_* = *f_I_th__*(*I_th_*) power relationship was further derived to be used in the creation of the final {*C*;*S_MN_*} and {*τ*; *S_MN_*} datasets in the next taken steps.

The adequacy of these final power trendlines and the statistical significance of the correlations were assessed identically to the {*A*; *B*} relationships derived directly from normalized experimental data, using the coefficient of determination *r*^2^ (squared value of the Pearson’s correlation coefficient) and a threshold of 0.01 on the *p*-value of the regression analysis, respectively. If *r*^2^ > 0.3 (i.e. *r* > 0.55) and *p* < 0.01, another power trendline *S_MN_* = *k_d_* · *A^d^* was fitted to the final dataset to describe the inverse relationship for {*S_MN_*; *A*} and used in the processing of the next-in-line property.

#### Normalized mathematical relationships between electrophysiological properties

When a final relationship with *S_MN_* was obtained for two MN properties *A* and *B* by the procedure described above, i.e., 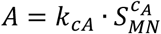 and 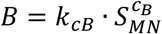, one of the expressions was mathematically inverted and a third empirical relationship was derived for the property pair {*A*; *B*}:

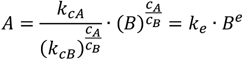

This procedure was applied to all possible {*A*; *B*} pairs in Table 1.

#### Validation of the normalized relationships

The normalized relationships were validated using a standard five-fold cross-validation procedure (Chollet, 2021). The data in each {*A*; *B*} global normalized dataset was initially randomized and therefore made independent from the studies constituting it. Each shuffled global dataset was then split into five non-overlapping complementary partitions, each containing 20% of the data. Four partitions including 80% of the data were taken to constitute a training set from which the normalized relationships were built as described previously. The latter equations were validated against the last data partition, that includes the remaining 20% of the data and which is called test set in the following. To perform this validation, the final size-related relationships 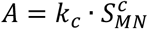 and relationships between electrophysiological properties *A* = *k_e_* · *B^e^* obtained with the training set were applied to the *S_MN_* or *B* data in each test set, yielding predicted values *A*. It was then assessed to what extent the mathematical relationships predicted the test data *A* by calculating the normalized maximum error (*nME*), the normalized root mean squared error (*nRMSE*) and the coefficient of determination 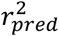 between predicted and control values of *A* in each test set. This process was repeated for a total of 5 times by permutating the five data partitions and creating five different pairs of one training set and one test set. The *nME, nRMSE* and 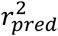 values were finally averaged across the five permutations. For each global dataset, the average 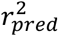 was compared to the 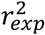 obtained from the power trendlines least-squared fitted to the control data in the log-log space. Once validated with this five-fold cross-validation method, the final normalized size-dependent relationships 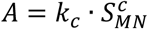 and relationships between electrophysiological properties *A* = *k_e_* · *B^e^* were computed for the complete global datasets.

#### Scaling of the normalized relationships

The 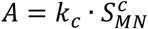 and *A* = *k_e_* · *B^e^* normalized relationships were finally scaled to typical cat property values in three steps. First, it was assessed from the literature the fold-range over which each MN property was to vary. *S_MN_* varied over a 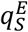-fold range, taken as the average across cat studies of the ratios of minimum and maximum values measured for *S_MN_:*

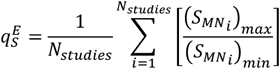

An experimental ratio 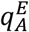 was similarly obtained for each electrophysiological property *A*. As 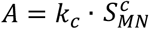, any electrophysiological property *A* could theoretically vary over a 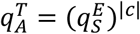-fold range, which was compared to 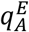 for consistency in the results. Then, a theoretical range [*A_min_*; *A_max_*] of values were derived for each electrophysiological property *A*. Defining 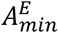 and 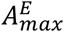 as the respective average of the minimum and maximum *A*-values across studies, it was enforced 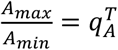 and 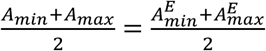 so that the [*A_min_*; *A_max_*] theoretical ranges were consistent with the normalized size-dependent relationships while best reproducing the experimental data from cat literature. A theoretical range for *S_MN_* [(*S_MN_*)_*min*_; (*S_MN_*)*_max_*] was similarly built over the 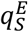-fold range previously derived. Finally, the intercept *k_c_* in the size-dependent relationships 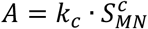 was scaled as:

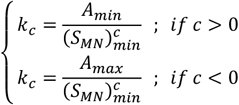

A similar approach was used to mathematically derive the intercept *k_e_* and scale the relationships *A* = *k_e_* · *B^e^* between electrophysiological properties.

#### Extrapolation to rats and mice

It was finally assessed whether the 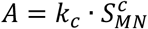 and *A* = *k_e_* · *B^e^* scaled relationships derived from cat data accurately predicted rat and mouse quantities reported in other experimental studies. These mathematical relationships were applied to the *S_MN_* or *B* data in each {*A*; *B*} global dataset of absolute values obtained for rats and mice independently. The accuracy in predicting the quantity *A* was assessed for each available dataset with the same three validation metrics used to validate the normalized power trend lines: the normalized maximum error (*nME*), the normalized root-mean-square error (*nRMSE*) and the coefficient of determination *r*^2^ between predicted and experimental values of A.

### Definitions of MN Size *S_MN_* and mU size *S_mU_*

#### MN Size *S_MN_*

The selected studies that performed both electrophysiological and morphometric measurements on the same MNs dominantly measured the MN soma diameter *D_soma_* as an index of MN size. Therefore, we chose *S_MN_* = *D_soma_* in the described methodology. In the literature on spinal MNs in adult mammals, the size of a MN *S_MN_* is also related to the measures of the somal cross-sectional area *CSA_soma_* (Gao et al., 2009; Friese et al., 2009; Deardorff et al., 2013; Mierzejewska-Krzyżowska et al., 2014; Dukkipati et al., 2018), soma surface area *S_soma_* (Ulfhake & Kellerth, 1984; Brandenburg et al., 2020), axonal diameter *D_axon_* (Cullheim, 1978), individual dendrite diameter *D_dendrite_* (Amendola & Durand, 2008; Carrascal et al., 2009; Mantilla et al., 2018), individual dendritic surface area *S_dendrite_* (Li et al., 2005; Obregón et al., 2009; Carrascal et al., 2009; Filipchuk & Durand, 2012; Kanjhan et al., 2015), total dendritic surface area (Ulfhake & Cullheim, 1988; Amendola & Durand, 2008; Brandenburg et al., 2020) or total MN surface area *S_neuron_* (Burke et al., 1982) defined as the summed soma and dendritic surface areas.

In the references (Zwaagstra & Kernell, 1981; Ulfhake & Kellerth, 1981), a linear correlation between *D_soma_* and the average diameter of the stem dendrites *D_dendrite_mean__* has been reported (*r* > 0.62, population size in {40; 82} cells). Moreover, a linear or quasi-linear correlation has been found (Cullheim et al., 1987; Ulfhake & Cullheim, 1988; Moschovakis et al., 1991; Prakash et al., 2000; Li et al., 2005; Obregón et al., 2009; Mantilla et al., 2018) between the stem dendrite diameter *D_dendrite_* and the membrane surface area of the corresponding dendritic tree *S_dendrite_* (*r* > 0.78, population size in [33; 342] dendrites). Therefore, from these studies, we can assume an approximate linear correlation between *D_soma_* and the average dendritic surface area *S_dendrite_mean__*. Moreover, the number of dendritic trees *N_dendrites_* per cell increases with increasing soma surface *S_soma_* (Brandenburg et al., 2018), and a linear correlation between *D_soma_* or *S_dendrites_* with *N_dendrites_* has been observed (Zwaagstra & Kernell, 1981; Ulfhake & Cullheim, 1988; Amendola & Durand, 2008) (*r* > 0.40, population size in {14; 32; 87} cells). It was therefore assumed that *D_soma_* and total dendritic surface area 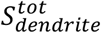 are linearly correlated. This assumption/approximation is consistent other conclusions of a linear correlations between *D_soma_* and 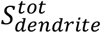 (Amendola & Durand, 2008; Brandenburg et al., 2020). Then, according to typical values of *D_soma_, S_soma_* and 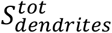 obtained from the studies previously cited, yielding 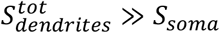, it was also assumed 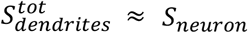. It is thus concluded that *D_soma_* is linearly related to total neuron membrane area *S_neuron_*, consistently with results by Burke et al. (1982) (*r* = 0.61, 57 cells).

For the above reasons and assumptions, the mathematical relationships 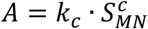 derived previously, with *S_MN_* = *D_soma_*, were extrapolated with a gain to relationships between *A* and *S_MN_* = *S_neuron_*, notably permitting the definition of surface specific resistance *R_m_* and capacitance *C_m_*. Following the same method as described above, theoretical ranges [*(S_neuron_*)*_min_*; (*S_neuron_*)*_max_*] were derived from additional morphometric studies on adult cat spinal alpha-motoneurons. The new relationships were extrapolated as 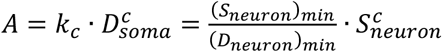.

It must however be highlighted that the conclusion of a linear correlation between *D_soma_* and *S_neuron_*, while plausible, is crude. Morphometric measurements of individual MNs are indeed difficult and suffer many limitations. MN staining, slice preparations and MN reconstruction and identification are complex experimental procedures requiring a large amount of work; the results from most of the cited studies thus rely on relatively small pools of investigated MNs. Most cited studies did not account for tissue shrinkage after dehydration, while some may have failed to assess the full dendritic trees from MN staining techniques (Brandenburg et al., 2020), thus underestimating dendritic membrane measurements. Moreover, before automated tools for image segmentation, landmark mapping and surface tracking (Amendola & Durand, 2008; Obregón et al., 2009; Mierzejewska-Krzyżowska et al., 2014) were available, morphometric measurements were performed from the manual reproduction of the cell outline under microscope, yielding important operator errors reported to be of the order of ~0.5*μm*, i.e. ~20% stem diameter. In these studies, morphologic quantities were besides obtained from geometrical approximations of the 2D MN shape, such as modelling the individual dendritic branches as one (Cullheim et al., 1987) or more (Prakash et al., 2000) equivalent cylinders for the derivation of *S_dendrite_*· Morphometric measurement approaches also varied between studies; for example, oval or circle shapes were best fitted by sight onto the soma outline and equivalent *D_soma_* quantities were derived either as the mean of measured maximum and minimum oval diameters or as the equivalent diameter of the fitted circle. In these studies, *CSA_soma_* and *S_soma_* were therefore derived by classical equations of circle and spheric surface areas, which contradicts the results by Mierzejewska-Krzyżowska et al. (2014) obtained from surface segmentation of a linear relationship between *D_soma_* and *CSA_soma_* (*r* = 0.94, 527 MNs). Finally, no correlation between soma size and *N_dendrites_* was found (Ulfhake & Kellerth, 1981; Cullheim et al., 1987). Therefore, the linear correlation between *D_soma_* and *S_neuron_* is plausible but crude and requires awareness of several important experimental limitations and inaccuracies. Conclusions and predictions involving *S_neuron_* should be treated with care in this study as these morphometric measurements lack the precision of the measures performed for the electrophysiological properties listed in Table 1.

#### Muscle unit size *S_mU_*

To enable future comparisons between MN and mU properties, we here assess potential indices of the mU size *S_mU_* suggested in the literature. The size of a mU (*S_mU_*) can be defined as the sum *CSA^tot^* of the cross-sectional areas CSAs of the innervated fibres composing the mU. *CSA^tot^* depends on the mU innervation ratio (*IR*) and on the mean CSA (*CSA^mean^*) of the innervated fibres: *S_mU_* = *CSA^tot^* = *IR* · *CSA^mean^*. *CSA^tot^* was measured in a few studies on cat and rat muscles, either indirectly by histochemical fibre profiling (Burke & Tsairis, 1973; Dum & Kennedy, 1980; Burke, 1981), or directly by glycogen depletion, periodic acid Schiff (PAS) staining and fibre counting (Burke et al., 1982; Rafuse et al., 1997). The mU tetanic force *F^tet^* is however more commonly measured in animals. As the fibre mean specific force *σ* is considered constant among the mUs of one muscle in animals (Lucas et al., 1987; Enoka, 1995), the popular equation *F^tet^* = *σ* · *IR*· *CSA^mean^* (Burke, 1981; Enoka, 1995) returns a linear correlation between *F^tet^* and *IR* · *CSA^mean^* = *S_mU_* in mammals. Experimental results (Burke & Tsairis, 1973; Bodine et al., 1987; Chamberlain & Lewis, 1989; Kanda & Hashizume, 1992; Rafuse et al., 1997; Hegedus et al., 2007) further show a linear correlation between *F^tet^* and *IR* and between *F^tet^* and *CSA^mean^*. Consequently, *F^tet^, IR* and *CSA^mean^* are measurable, consistent, and linearly related measures of *S_mU_* in animals, as summarized in Table 2.

**Table 2.** Measurable indices of MN and mU sizes in mammals. *S_MN_* and *S_mU_* are conceptual parameters which are adequately described by the measurable and linearly inter-related quantities reported in this table.

| MN size ($S_{MN}$) | mU size ($S_{mU}$) |
| --- | --- |
| $S_{neuron}$ | $F^{tet}$ |
| $D_{soma}$ | $IR$ |
| | $CSA^{mean}$ |
| | $CSA^{tot}$ |

#### Relationships between MN and mU properties

To assess whether the size-dependent relationships 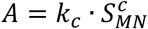 derived in this study were in accordance with Henneman’s size principle of motor unit recruitment, we identified a set of experimental studies that concurrently measured a MN property *B_MN_* (Table 1) and a muscle unit (mU) property *A_mU_* for the same MU. The normalized global datasets obtained for the pairs {*A_mU_; B_MN_*} were fitted with power trendlines, as previously described for MN properties, yielding 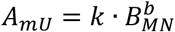 relationships. Using both the definition of *S_mU_* (Table 2) and the 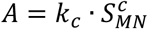 relationships derived previously, the 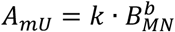 relationships were then mathematically transformed into 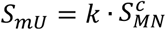 relationships. If all *c*-values obtained from different {*A_mU_; B_MN_*} pairs were of the same sign, it was concluded that mU and MN sizes were monotonically related. Considering the limited data available obtained from the literature, data obtained for MNs innervating different hindlimb muscles were merged. Also, cat, rat and mouse studies were processed independently but the resulting *c*-values were compared without regards to the related species.

## Results

We respectively identified 19, 6 and 4 studies respecting our desired criteria on cats, rats and mice that reported processable experimental data for the morphometric and electrophysiological properties listed in Table 1. Additional publications including some from the past 10 years were identified but could not be included in this work as no processable data could be recovered. An additional 14 studies were found to perform concurrent MN and mU measurements on individual motor units. From the selected cat studies, the 17 pairs of MN properties and the 8 pairs of one MN and one mU property represented in the bubble diagram of Figure 2A were investigated.

**Figure 2.**
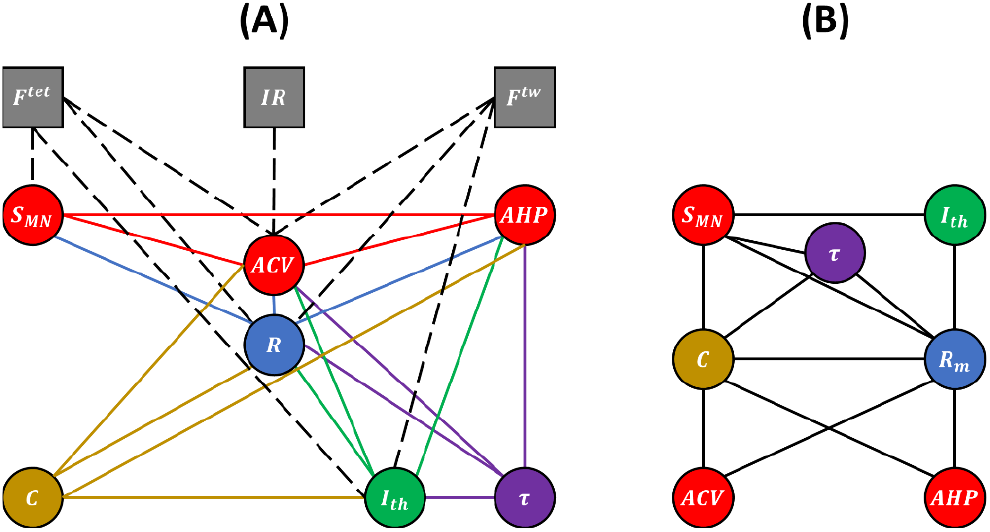
Experimental (A) and unknown (B) relations between MN and mU properties. (A) Bubble diagram representing the pairs of MN and/or mU properties that could be investigated in this study from the results provided by the 40 studies identified in our web search. MN and mU properties are represented in circle and square bubbles, respectively. Relationships between MN properties are represented by coloured connecting lines; the colours red, blue, green, yellow, and purple are consistent with the order *ACV, AHP, R, I_th_, C, τ* in which the pairs were investigated (see Table 3 for mathematical relationships). Relationships between one MN and one mU property are represented by black dashed lines. (B) Bubble diagram representing the mathematical relationships proposed in this study between pairs of MN properties for which no concurrent experimental data has been measured to date.

### Relationships between MN properties

#### Global datasets and data variability

The experimental data retrieved from the selected studies for the 17 pairs of MN properties drawn in Figure 2A were merged into 17 normalized global datasets plotted in Figure 3. Eight global datasets ({*ACV; S_MN_*}, {*AHP; ACV*}, {*R; S_MN_*}, {*R; ACV*}, {*R; AHP*}, {*I_th_*; *R*}, {*I_th_*; *ACV*}, { *τ*; *R*}) included the data from two or more experimental studies.

**Figure 3.**
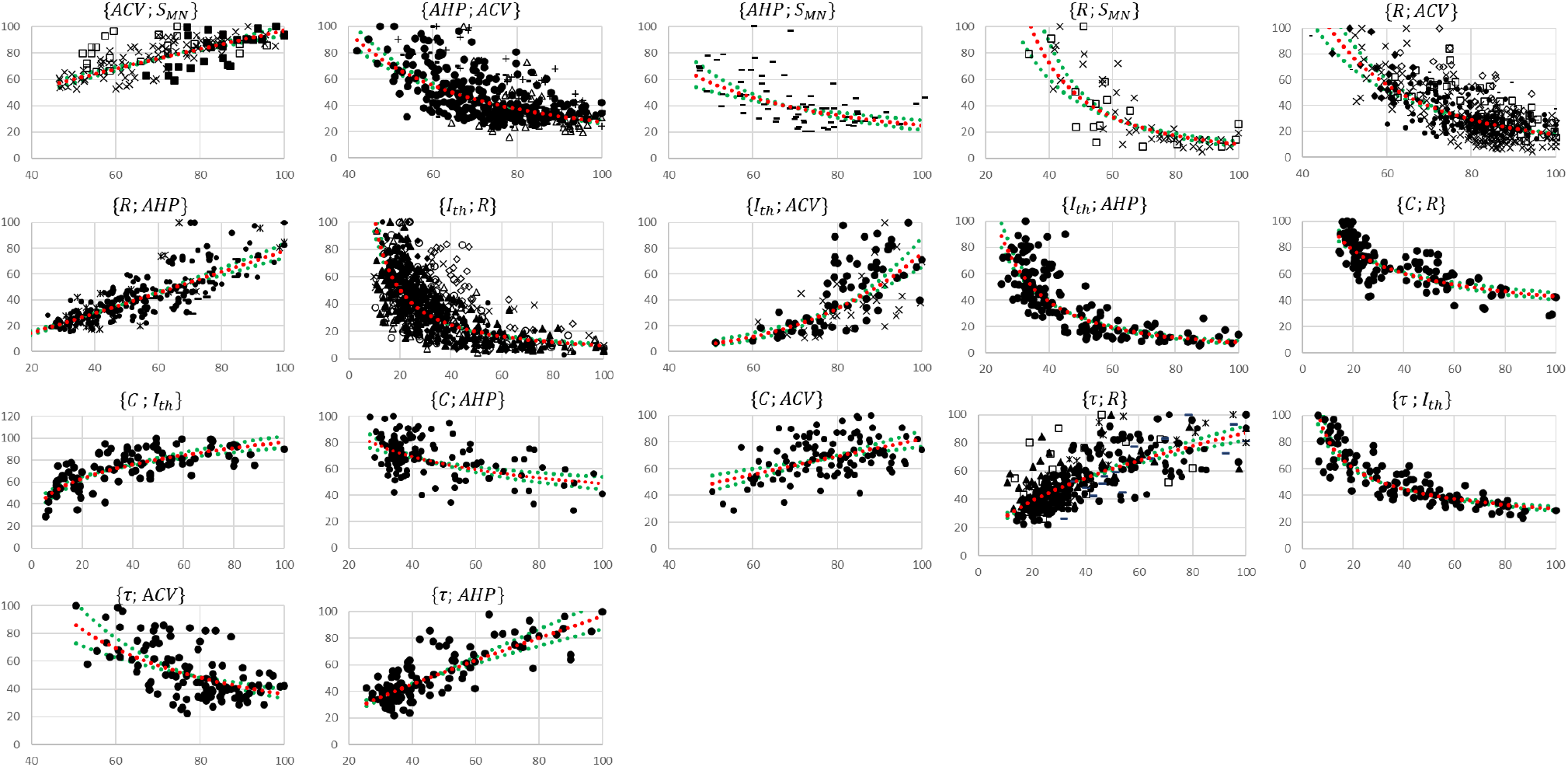
Normalized global datasets. These are obtained from the 19 studies reporting cat data that measured and investigated the 17 pairs of MN properties reported in Figure 2A. For each {*A;B*} pair, the property *A* is read on the y axis and *B* on the x axis. For information, power trendlines *A* = *k* · *B^a^* (red dotted curves) are fitted to the data of each dataset and reported in Table 3. The 95% confidence interval of the regression is also displayed for each dataset (green dotted lines). The studies are identified with the following symbols: • (Gustafsson, 1979; Gustafsson & Pinter, 1984a; Gustafsson & Pinter, 1984b), ○ (Munson et al., 1986), ▲ (Zengel et al., 1985), Δ (Foehring et al., 1987), ■ (Cullheim, 1978), □ (Burke, 1968; Burke & Ten Bruggencate, 1971; Burke et al., 1982), ◆ (Krawitz et al., 2001), ◇ (Fleshman et al., 1981), + (Eccles et al., 1958), X (Kernell, 1966; Kernell & Zwaagstra, 1981; Kernell & Monster, 1981), - (Zwaagstra & Kernell, 1980), – (Sasaki, 1991), ✷ (Pinter & Vanden Noven, 1989). The axes are given in % of the maximum retrieved values in the studies consistently with the Methods section.

These studies reported in each global dataset similar normalized property distributions, as visually displayed in the frequency histograms in Figure FSM1 in Supplementary Material. This was confirmed by the *mean_g_* ± *sd_g_* calculations described in the Methods of the four metrics range, mean, CoV, and 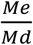 displayed as error bars in Figure FSM2 in Supplementary Material. Out of the 64 *mean_g_+/-sd_g_* calculations (four metrics for 16 property distributions of the 8 global datasets), 55 (86%) returned *sd_g_* < 10% and 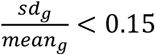. Otherwise, the distributions of only three, two and four properties showed sensibly higher inter-study variability in the global datasets for the range, mean and CoV metrics respectively, however still verifying *sd_g_* < 15% and 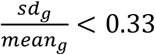. Additional details are provided in Supplementary Material (Section SM1 and Figure FSM2). Then, as displayed in Figure 3 and reported in Table 3, power trendlines and their 95% confidence interval were fitted to the global datasets. All 17 trendlines were statistically significant and adequately represented by a power model *A* = *k_a_* · *B^a^* (*p*-value < 10^−5^ and *r*^2^ ∈ [0.34; 0.72] for 15 datasets; *p*-value < 10^−5^ and *r*^2^ ∈ {0.24; 0.17} for the {*C*; *AHP*} and {*C*; *CV*} datasets respectively). As displayed in green dotted lines in Figure 3, the confidence interval remained narrow for all datasets with the value of power *a* varying less than 0.4, except for the {*I_th_;ACV*} dataset, suggesting a low inter-study variability in associating the distributions of two properties. As displayed in Figure FSM3 in Supplementary Material, the selected studies however reported datasets of different sizes. Yet, out of the 35 experimental datasets constituting the 8 global datasets identified previously, only one and seven experimental datasets were identified to respectively constitute more than 50% and less than 10% of the global dataset they were included in. Therefore, despite a general imbalance towards the over-appearance of the work from Gustafsson & Pinter (1984ab) and a tendency towards overlooking some small datasets (Kernell, 1966; Burke & Ten Bruggencate, 1971; Sasaki, 1991), the remaining studies all played a significant role in the procedure to constitute the final datasets. Therefore, the inter-study data variability was globally low, and the data distributions reported in the experimental studies were confidently merged into the global datasets plotted in Figure 3. It is worth noting that three research groups provided 40%, 27% and 15% of the 2717 data points describing the *S_MN_, ACV, AHP, R, I_th_* and *τ* property distributions in this study, while the C-related data was provided by one only research group, leading to potential group-specific methodological bias.

**Table 3.**
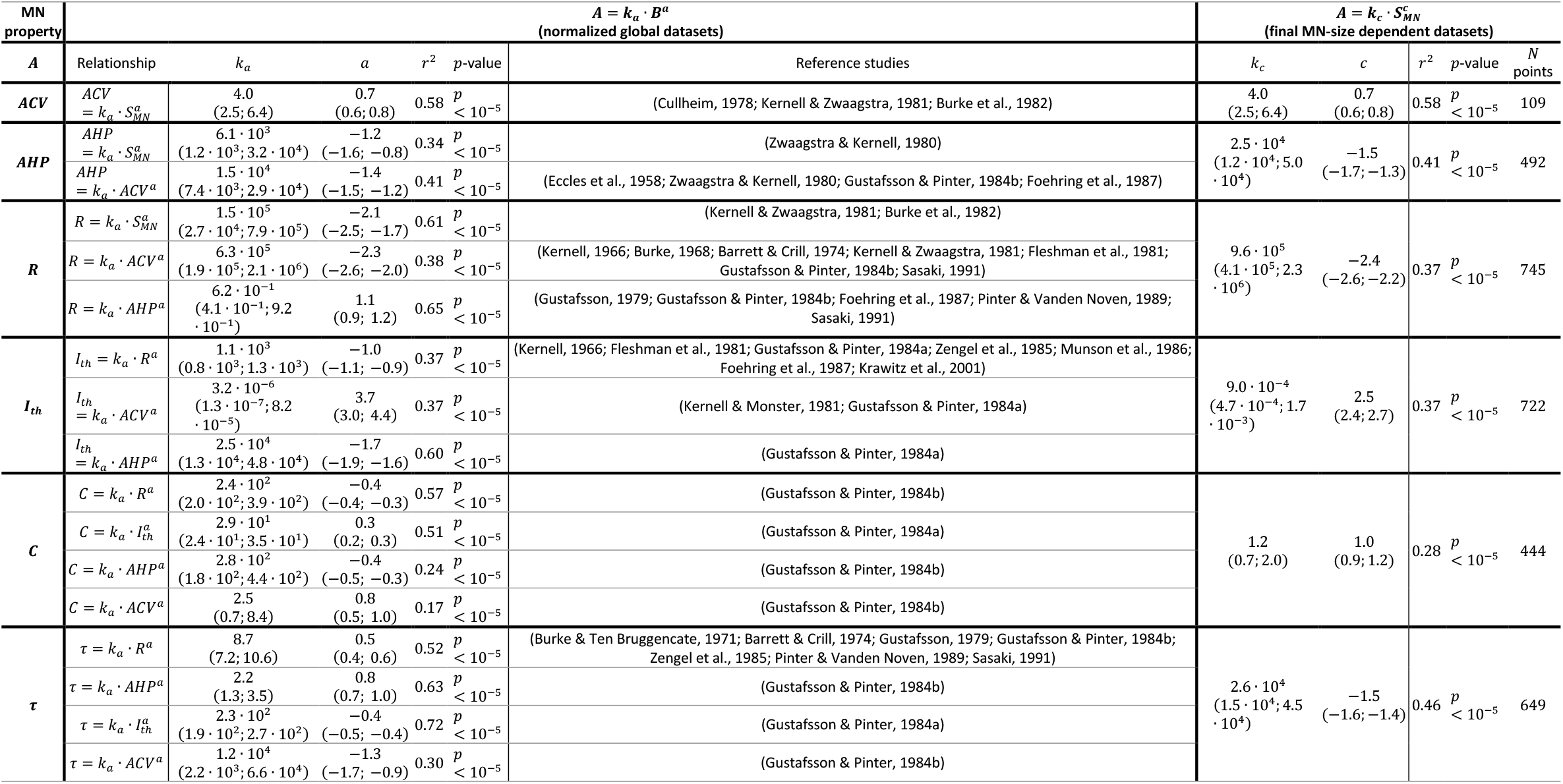
Fitted experimental data of pairs of MN properties and subsequent normalized final size-related relationships. For information, the *r*^2^, *p*-value and the equation *A* = *k_a_* · *B^a^* are reported for each fitted global dataset. The normalized MN-size dependent relationships 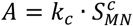 are mathematically derived from the transformation of the global datasets and from the power trendline fitting of the final datasets (N data points) as described in the Methods. The minimum and maximum values of *k_a_, k_c_, a* and *c* defining the 95% confidence interval of the regression are also reported in parenthesis for each global and final dataset. The *r*^2^ values reported in this table are consistent with the *r*^2^ values obtained when directly fitting the normalized experimental datasets with power regressions (see Table TSM1 in Supplementary Material).

| MN property | $A = k_a \cdot B^a$<br>(normalized global datasets) | | | | | $A = k_c \cdot S_{MN}^c$<br>(final MN-size dependent datasets) | | | | | |
| --- | --- | --- | --- | --- | --- | --- | --- | --- | --- | --- | --- |
| A | Relationship | $k_a$ | $a$ | $r^2$ | p-value | Reference studies | $k_c$ | $c$ | $r^2$ | p-value | $N$<br>points |
| ACV | $ACV = k_a \cdot S_{MN}^a$ | 4.0<br>(2.5; 6.4) | 0.7<br>(0.6; 0.8) | 0.58 | $p < 10^{-5}$ | (Cullheim, 1978; Kernell & Zwaagstra, 1981; Burke et al., 1982) | 4.0<br>(2.5; 6.4) | 0.7<br>(0.6; 0.8) | 0.58 | $p < 10^{-5}$ | 109 |
| AHP | $AHP = k_a \cdot S_{MN}^a$ | $6.1 \cdot 10^3$<br>($1.2 \cdot 10^3$; $3.2 \cdot 10^4$) | -1.2<br>(-1.6; -0.8) | 0.34 | $p < 10^{-5}$ | (Zwaagstra & Kernell, 1980) | $2.5 \cdot 10^4$<br>($1.2 \cdot 10^4$; $5.0 \cdot 10^4$) | -1.5<br>(-1.7; -1.3) | 0.41 | $p < 10^{-5}$ | 492 |
| | $AHP = k_a \cdot ACV^a$ | $1.5 \cdot 10^4$<br>($7.4 \cdot 10^3$; $2.9 \cdot 10^4$) | -1.4<br>(-1.5; -1.2) | 0.41 | $p < 10^{-5}$ | (Eccles et al., 1958; Zwaagstra & Kernell, 1980; Gustafsson & Pinter, 1984b; Foehring et al., 1987) | | | | | |
| R | $R = k_a \cdot S_{MN}^a$ | $1.5 \cdot 10^5$<br>($2.7 \cdot 10^4$; $7.9 \cdot 10^5$) | -2.1<br>(-2.5; -1.7) | 0.61 | $p < 10^{-5}$ | (Kernell & Zwaagstra, 1981; Burke et al., 1982) | $9.6 \cdot 10^5$<br>($4.1 \cdot 10^5$; $2.3 \cdot 10^6$) | -2.4<br>(-2.6; -2.2) | 0.37 | $p < 10^{-5}$ | 745 |
| | $R = k_a \cdot ACV^a$ | $6.3 \cdot 10^5$<br>($1.9 \cdot 10^5$; $2.1 \cdot 10^6$) | -2.3<br>(-2.6; -2.0) | 0.38 | $p < 10^{-5}$ | (Kernell, 1966; Burke, 1968; Barrett & Crill, 1974; Kernell & Zwaagstra, 1981; Fleshman et al., 1981; Gustafsson & Pinter, 1984b; Sasaki, 1991) | | | | | |
| | $R = k_a \cdot AHP^a$ | $6.2 \cdot 10^{-1}$<br>($4.1 \cdot 10^{-1}$; $9.2 \cdot 10^{-1}$) | 1.1<br>(0.9; 1.2) | 0.65 | $p < 10^{-5}$ | (Gustafsson, 1979; Gustafsson & Pinter, 1984b; Foehring et al., 1987; Pinter & Vanden Noven, 1989; Sasaki, 1991) | | | | | |
| $I_{th}$ | $I_{th} = k_a \cdot R^a$ | $1.1 \cdot 10^3$<br>($0.8 \cdot 10^3$; $1.3 \cdot 10^3$) | -1.0<br>(-1.1; -0.9) | 0.37 | $p < 10^{-5}$ | (Kernell, 1966; Fleshman et al., 1981; Gustafsson & Pinter, 1984a; Zengel et al., 1985; Munson et al., 1986; Foehring et al., 1987; Krawitz et al., 2001) | $9.0 \cdot 10^{-4}$<br>($4.7 \cdot 10^{-4}$; $1.7 \cdot 10^{-3}$) | 2.5<br>(2.4; 2.7) | 0.37 | $p < 10^{-5}$ | 722 |
| | $I_{th} = k_a \cdot ACV^a$ | $3.2 \cdot 10^{-6}$<br>($1.3 \cdot 10^{-7}$; $8.2 \cdot 10^{-5}$) | 3.7<br>(3.0; 4.4) | 0.37 | $p < 10^{-5}$ | (Kernell & Monster, 1981; Gustafsson & Pinter, 1984a) | | | | | |
| | $I_{th} = k_a \cdot AHP^a$ | $2.5 \cdot 10^4$<br>($1.3 \cdot 10^4$; $4.8 \cdot 10^4$) | -1.7<br>(-1.9; -1.6) | 0.60 | $p < 10^{-5}$ | (Gustafsson & Pinter, 1984a) | | | | | |
| C | $C = k_a \cdot R^a$ | $2.4 \cdot 10^2$<br>($2.0 \cdot 10^2$; $3.9 \cdot 10^2$) | -0.4<br>(-0.4; -0.3) | 0.57 | $p < 10^{-5}$ | (Gustafsson & Pinter, 1984b) | 1.2<br>(0.7; 2.0) | 1.0<br>(0.9; 1.2) | 0.28 | $p < 10^{-5}$ | 444 |
| | $C = k_a \cdot I_{th}^a$ | $2.9 \cdot 10^1$<br>($2.4 \cdot 10^1$; $3.5 \cdot 10^1$) | 0.3<br>(0.2; 0.3) | 0.51 | $p < 10^{-5}$ | (Gustafsson & Pinter, 1984a) | | | | | |
| | $C = k_a \cdot AHP^a$ | $2.8 \cdot 10^2$<br>($1.8 \cdot 10^2$; $4.4 \cdot 10^2$) | -0.4<br>(-0.5; -0.3) | 0.24 | $p < 10^{-5}$ | (Gustafsson & Pinter, 1984b) | | | | | |
| | $C = k_a \cdot ACV^a$ | 2.5<br>(0.7; 8.4) | 0.8<br>(0.5; 1.0) | 0.17 | $p < 10^{-5}$ | (Gustafsson & Pinter, 1984b) | | | | | |
| $\tau$ | $\tau = k_a \cdot R^a$ | 8.7<br>(7.2; 10.6) | 0.5<br>(0.4; 0.6) | 0.52 | $p < 10^{-5}$ | (Burke & Ten Bruggencate, 1971; Barrett & Crill, 1974; Gustafsson, 1979; Gustafsson & Pinter, 1984b; Zengel et al., 1985; Pinter & Vanden Noven, 1989; Sasaki, 1991) | $2.6 \cdot 10^4$<br>($1.5 \cdot 10^4$; $4.5 \cdot 10^4$) | -1.5<br>(-1.6; -1.4) | 0.46 | $p < 10^{-5}$ | 649 |
| | $\tau = k_a \cdot AHP^a$ | 2.2<br>(1.3; 3.5) | 0.8<br>(0.7; 1.0) | 0.63 | $p < 10^{-5}$ | (Gustafsson & Pinter, 1984b) | | | | | |
| | $\tau = k_a \cdot I_{th}^a$ | $2.3 \cdot 10^2$<br>($1.9 \cdot 10^2$; $2.7 \cdot 10^2$) | -0.4<br>(-0.5; -0.4) | 0.72 | $p < 10^{-5}$ | (Gustafsson & Pinter, 1984a) | | | | | |
| | $\tau = k_a \cdot ACV^a$ | $1.2 \cdot 10^4$<br>($2.2 \cdot 10^3$; $6.6 \cdot 10^4$) | -1.3<br>(-1.7; -0.9) | 0.30 | $p < 10^{-5}$ | (Gustafsson & Pinter, 1984b) | | | | | |

The merged property distributions obtained in the 17 global datasets showed low variability between global datasets, as displayed in Figure FSM4 in Supplementary Material. Indeed, *sd_G_* < 10% and 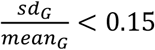 was obtained for all four metrics for all the investigated properties across the global datasets they appeared in. Therefore, the property distributions were similar enough between global datasets to apply the process displayed in Figure 1 and derive the final size-dependent datasets.

#### Size-dependent normalized relationships

The 17 normalized global datasets were processed according to the procedure described in Figure 1 to derive normalized mathematical relationships between the electrophysiological properties listed in Table 1 and MN size *S_MN_*. First, the normalized size-dependent relationship 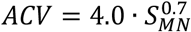, reported in Table 3, was derived from the trendline fitting of the normalized {*ACV; S_MN_*} global dataset, which was obtained from 3 studies (Cullheim, 1978; Kernell & Zwaagstra, 1981; Burke et al., 1982) and is represented in the upper-left panel in Figure 3. This resulted in a statistically significant relationship (*r*^2^ = 0.58, *p*-value< 10^−5^). A statistically significant (*r*^2^ = 0.59, *p*-value< 10^−5^) inverse relationship *S_MN_ = k_d_* · *ACV^d^* was also fitted to this dataset. Then, the {*AHP; ACV*} dataset, represented in the second panel upper row in Figure 3 and obtained from 4 studies (Eccles et al., 1958; Zwaagstra & Kernell, 1980; Gustafsson & Pinter, 1984b; Foehring et al., 1987) was transformed into a new {*AHP; S_MN_*}_1_ dataset by applying the priory derived *S_MN_* = *k_d_* · *ACV^d^* relationship to the list of *ACV*-values. This transformed {*AHP; S_MN_*}_1_ dataset was then merged with the {*AHP; S_MN_}*_2_ dataset (third panel upper row in Figure 2) obtained from (Zwaagstra & Kernell, 1980), yielding the final {*AHP; S_MN_}_f_* dataset of *N* = 492 data points shown in Figure 4, second panel. A statistically significant (*r*^2^ = 0.58, *p*-value< 10^−5^) power trendline 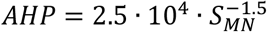 was fitted to the {*AHP; S_MN_}_f_* dataset and is reported in Table 3. As before, a statistically significant (*r*^2^ = 0.38, *p*-value < 10^−5^) inverse relationship *S_MN_* = *k_d_* · *AHP^d^* was also fitted to this dataset for future use. A similar procedure was applied to derive the normalized final relationships between {*R, I_th_, C, τ*} and *S_MN_* reported in the last column of Table 3. A statistically significant (*p*-value< 10^−5^) correlation was obtained between each electrophysiological property {*ACV,AHP,R,I_th_,C,τ*} and *S_MN_* as reported in Table 3. With *r*^2^ ∈ [0.28; 0.58], it was obtained that {*I_th_,C,CV*} and {*R,AHP,τ*} respectively increased and decreased with increasing MN sizes *S_MN_*, with slopes reported in Table 3 and plotted in Figure 5.

**Figure 4.**
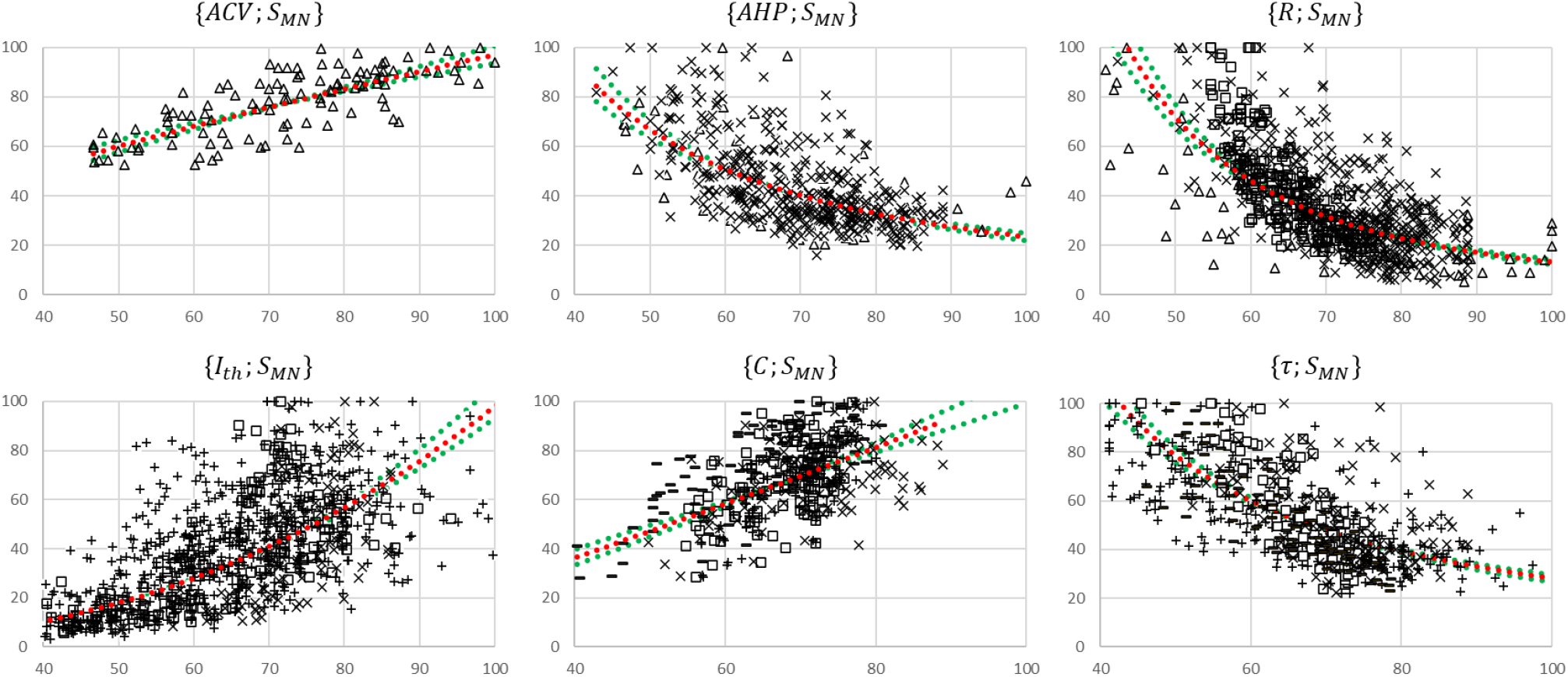
Normalized MN size-related final datasets. These are obtained from the 19 studies reporting cat data that concurrently measured at least two of the morphometric and electrophysiological properties listed in Table 1. For each {*A*; *S_MN_*} pair, the property *A* is read on the y axis and *S_MN_* on the x axis. The power trendlines 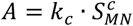 (red dotted curves) are fitted to each dataset and are reported in Table 3. The 95% confidence interval of the regression is also displayed for each dataset (green dotted lines). For each {*A*; *S_MN_*} plot, the constitutive sub-datasets {*A*; *S_MN_*} that were obtained from different global {*A;B*} datasets are specified with the following symbols identifying the property 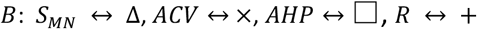, and *I_th_* ↔ –.

**Figure 5.**
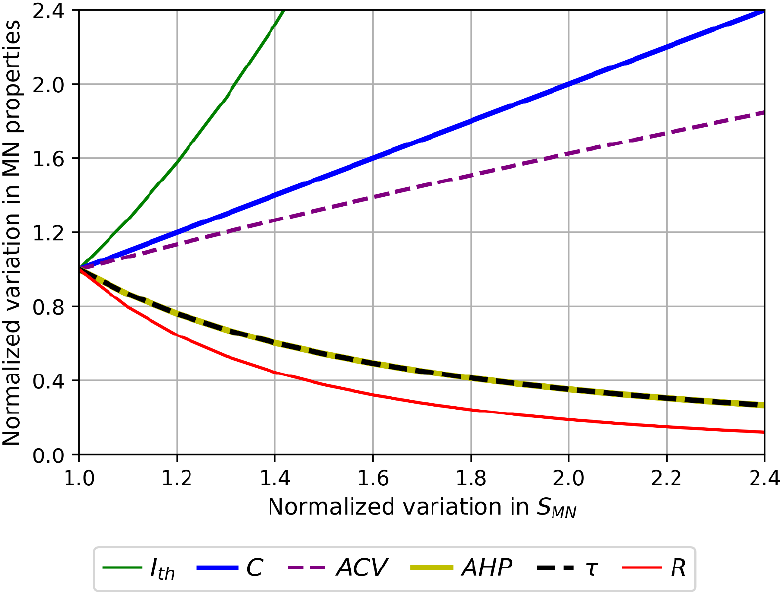
Normalized size-dependent behaviour of the MN properties *ACV,AHP,R,I_th_, C* and *τ*. For displaying purposes, the MN properties are plotted in arbitrary units as power functions (intercept *k* = 1) of 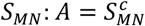 according to Table 3. The larger the MN size, the larger *ACV, C* and *I_th_* in the order of increasing slopes, and the lower *AHP, τ* and *R* in the order of increasing slopes.

#### Normalized mathematical relationships between electrophysiological properties

The final size-dependent relationships reported in Table 3 were mathematically combined to yield normalized relationships between all electrophysiological properties listed in Table 1. As an example, 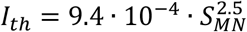 and 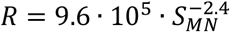 were obtained in Table 3. When mathematically combined (with the latter mathematically inverted as *S_MN_* = 2.9 · 10^2^ · *R*^−0.4^), these relationships yielded the normalized relationship between rheobase and input resistance *I_th_* = 1.5 · 10^3^ · *R^−1^*. All normalized relationships between MN electrophysiological properties hence obtained are provided in Supplementary Material.

#### Validation of the normalized relationships

The normalized relationships reported in Table TSM2 were obtained from the complete global datasets represented in Figure 3. A five-fold cross-validation of these relationships was performed, as described in the Methods section, and assessed with calculations of the *nME,nRMSE* and 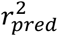 validation metrics averaged across the five permutations. As displayed in Figure 6, the normalized relationships obtained from the 17 training datasets predicted, in average across the five permutations, the experimental data from the test datasets with average *nME* between 52% and 300%, *nRMSE* between 13% and 24%, and coefficients of determination 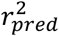 between 0.20 and 0.74 and of the same order of magnitude as the experimental 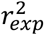 values.

**Figure 6.**
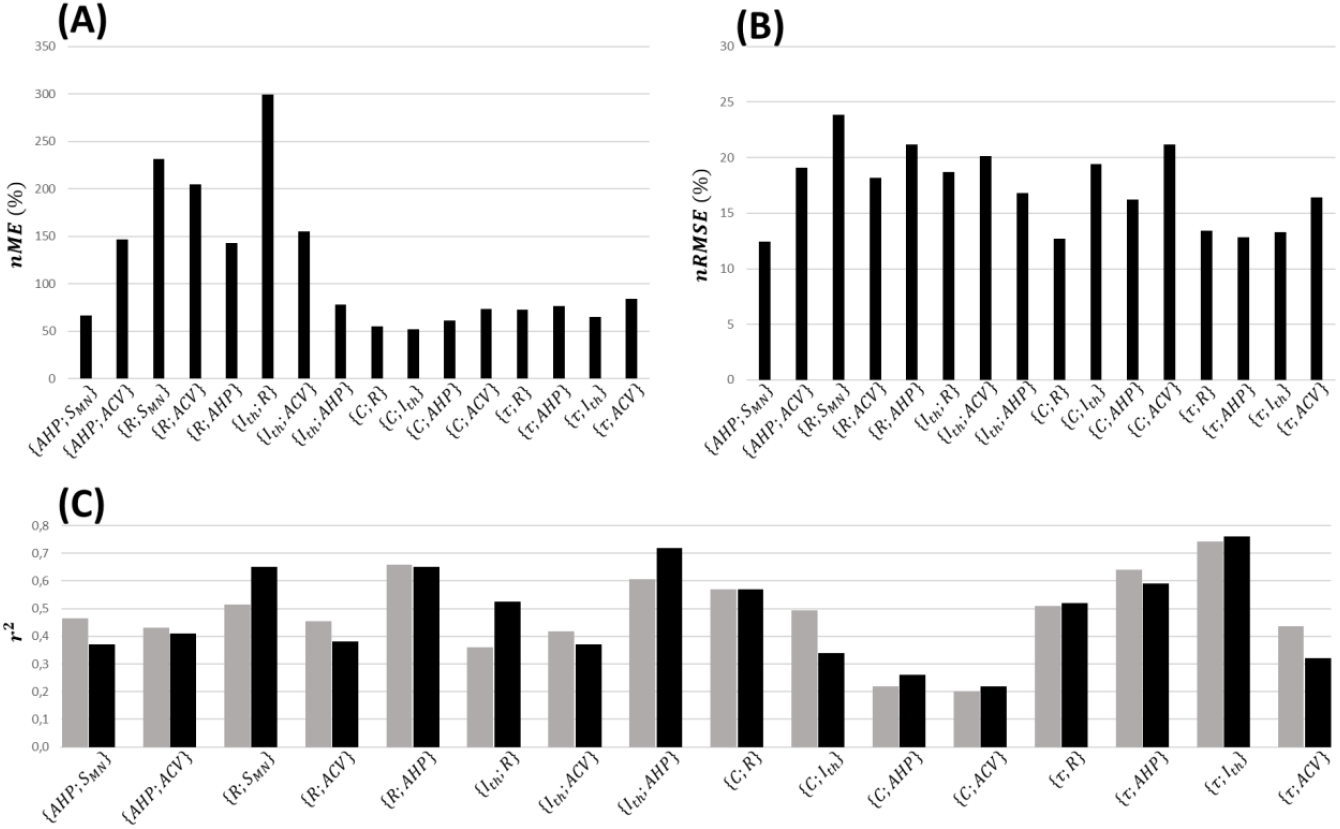
Five-fold cross-validation of the normalized mathematical relationships. Here are reported for each dataset the average values across the five permutations of (A) the normalized maximum error (*nME*), (B) the normalized root-mean-square error (*nRMSE*), and (C) coefficient of determination (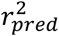, grey bars) which is compared with the coefficient of determination (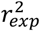, black bars) of the power trendline fitted to the log-log transformation of global experimental datasets directly.

#### Scaling of the normalized relationships

The normalized relationships provided in Supplementary Material were finally scaled to typical cat data obtained from the literature following the procedure described in the Methods section. *D_soma_* and *S_neuron_* were found to vary in cats over an average 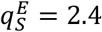-fold range according to a review of the morphometric studies reported in the two first lines of Table TSM3. 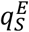 may however be underestimated for reasons discussed in Supplementary Material SM2.Then, the empirical 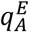 and theoretical 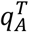 ratios, defined in the Methods, were calculated for each MN electrophysiological property *A* and are reported in Table TSM4. For example, MN resistance *R* was found to vary over an average 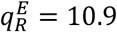-fold range in a MN pool according to the literature, while the theoretical fold range 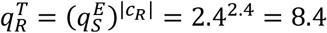 was obtained from the 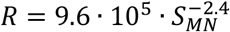 normalized relationship previously derived when a 2.4-fold range is set for *S_MN_*. As shown in Table TSM4, 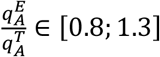 for the MN properties *ACV, AHP, C* and *τ*, while the theoretical ranges for *R* and *I_th_* span over a narrower domain than expected from experimental results (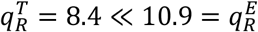 and 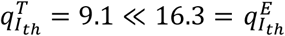). This suggests that the range of variation of *R* and *I_th_* is not entirely explained by the variation in MN size *S_MN_*, and that MN excitability is not only determined by *S_MN_*, as reviewed in Powers & Binder (2001).

The normalized relationships were finally scaled using the theoretical ranges reported in the last column of Table TSM4. Taking the {*I_th_;R*} pair as example, as *I_th_* = 1.4 · 10^3^ · *R*^−1^ (normalized relationship derived previously and provided in Supplementary Material), *R* ∈ [0.5; 4.0] · 10^6^*Ω* and *I_th_* ∈ [3.9; 35.0] · 10^−9^*A* (Table TSM4), it was directly obtained from the Methods that 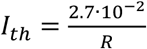 in SI base units. A similar approach yielded the final mathematical relationships reported in Table 4 between all MN electrophysiological and morphometric properties *ACV, AHP, R, I_th_, C, τ* and *S_MN_*. All constants and relationships are given in SI base units.

**Table 4.** Mathematical empirical cat relationships between the MN properties *S_neuron_, D_soma_, R, R_m_, C, τ, I_th_, AHP* and *ACV*. Each column provides the relationships between one and the eight other MN properties. If one property is known, the complete MN profile can be reconstructed by using the pertinent line in this table. All constants and properties are provided in SI base units (meters, seconds, ohms, farads and ampers). The relationships involving *R_m_* were obtained from theoretical relationships involved in Rall’s cable theory (See Discussion).

| | $S_{neuron}[m^2]$ | $D_{soma}[m]$ | $R[\Omega]$ | $R_m[\Omega \cdot m^2]$ | $C[F]$ | $\tau[s]$ | $I_{th}[A]$ | $AHP[s]$ | $ACV[m \cdot s^{-1}]$ |
| --- | --- | --- | --- | --- | --- | --- | --- | --- | --- |
| $S_{neuron}[m^2]$ | | $D_{soma} = 1.8 \cdot 10^2 \cdot S_{neuron}$ | $R = \frac{1.7 \cdot 10^{-10}}{S_{neuron}^{2.4}}$ | $R_m = \frac{1.0 \cdot 10^{-10}}{S_{neuron}^{1.4}}$ | $C = 1.3 \cdot 10^{-2} \cdot S_{neuron}$ | $\tau = \frac{1.0 \cdot 10^{-12}}{S_{neuron}^{1.5}}$ | $I_{th} = 3.8 \cdot 10^8 \cdot S_{neuron}^{2.5}$ | $AHP = \frac{1.0 \cdot 10^{-11}}{S_{neuron}^{1.5}}$ | $ACV = 3.0 \cdot 10^6 \cdot S_{neuron}^{0.7}$ |
| $D_{soma}[m]$ | $S_{neuron} = 5.5 \cdot 10^{-3} \cdot D_{soma}$ | | $R = \frac{5.1 \cdot 10^{-5}}{D_{soma}^{2.4}}$ | $R_m = \frac{1.7 \cdot 10^{-7}}{D_{soma}^{1.4}}$ | $C = 7.9 \cdot 10^{-5} \cdot D_{soma}$ | $\tau = \frac{2.3 \cdot 10^{-9}}{D_{soma}^{1.5}}$ | $I_{th} = 7.8 \cdot 10^2 \cdot D_{soma}^{2.5}$ | $AHP = \frac{2.7 \cdot 10^{-8}}{D_{soma}^{1.5}}$ | $ACV = 8.1 \cdot 10^4 \cdot D_{soma}^{0.7}$ |
| $R[\Omega]$ | $S_{neuron} = \frac{9.5 \cdot 10^{-5}}{R^{0.4}}$ | $D_{soma} = \frac{1.7 \cdot 10^{-2}}{R^{0.4}}$ | | $R_m = 5.7 \cdot 10^{-5} \cdot R^{0.6}$ | $C = \frac{1.5 \cdot 10^{-6}}{R^{0.4}}$ | $\tau = 9.6 \cdot 10^{-7} \cdot R^{0.6}$ | $I_{th} = \frac{2.7 \cdot 10^{-2}}{R}$ | $AHP = 1.3 \cdot 10^{-5} \cdot R^{0.6}$ | $ACV = \frac{4.9 \cdot 10^3}{R^{0.3}}$ |
| $R_m[\Omega \cdot m^2]$ | $S_{neuron} = \frac{1.5 \cdot 10^{-7}}{R_m^{0.7}}$ | $D_{soma} = \frac{2.6 \cdot 10^{-5}}{R_m^{0.7}}$ | $R = 6.8 \cdot 10^6 \cdot R_m^{1.7}$ | | $C = \frac{1.8 \cdot 10^{-9}}{R_m^{0.7}}$ | $\tau = 1.4 \cdot 10^{-2} \cdot R_m$ | $I_{th} = \frac{2.2 \cdot 10^{-9}}{R_m^{1.8}}$ | $AHP = 2.2 \cdot 10^{-1} \cdot R_m^{1.1}$ | $ACV = \frac{5.4 \cdot 10^1}{R_m^{0.5}}$ |
| $C[F]$ | $S_{neuron} = 1.2 \cdot 10^2 \cdot C$ | $D_{soma} = 2.1 \cdot 10^4 \cdot C$ | $R = \frac{1.5 \cdot 10^{-15}}{C^{2.5}}$ | $R_m = \frac{1.7 \cdot 10^{-13}}{C^{1.5}}$ | | $\tau = \frac{8.6 \cdot 10^{-16}}{C^{1.5}}$ | $I_{th} = 6.5 \cdot 10^{13} \cdot C^{2.6}$ | $AHP = \frac{7.7 \cdot 10^{-15}}{C^{1.6}}$ | $ACV = 8.2 \cdot 10^7 \cdot C^{0.7}$ |
| $\tau[s]$ | $S_{neuron} = \frac{8.4 \cdot 10^{-9}}{\tau^{0.7}}$ | $D_{soma} = \frac{1.5 \cdot 10^{-6}}{\tau^{0.7}}$ | $R = 7.1 \cdot 10^9 \cdot \tau^{1.6}$ | $R_m = 3.6 \cdot 10^1 \cdot \tau$ | $C = \frac{1.6 \cdot 10^{-10}}{\tau^{0.7}}$ | | $I_{th} = \frac{1.6 \cdot 10^{-12}}{\tau^{1.7}}$ | $AHP = 1.7 \cdot 10^1 \cdot \tau$ | $ACV = \frac{7.5}{\tau^{0.5}}$ |
| $I_{th}[A]$ | $S_{neuron} = 4.0 \cdot 10^{-4} \cdot I_{th}^{0.4}$ | $D_{soma} = 7.1 \cdot 10^{-2} \cdot I_{th}^{0.4}$ | $R = \frac{3.1 \cdot 10^{-2}}{I_{th}}$ | $R_m = \frac{7.4 \cdot 10^{-6}}{I_{th}^{0.6}}$ | $C = 5.9 \cdot 10^{-6} \cdot I_{th}^{0.4}$ | $\tau = \frac{1.2 \cdot 10^{-7}}{I_{th}^{0.6}}$ | | $AHP = \frac{1.5 \cdot 10^{-6}}{I_{th}^{0.6}}$ | $ACV = 1.3 \cdot 10^4 \cdot I_{th}^{0.3}$ |
| $AHP[s]$ | $S_{neuron} = \frac{5.5 \cdot 10^{-8}}{AHP^{0.7}}$ | $D_{soma} = \frac{9.8 \cdot 10^{-6}}{AHP^{0.7}}$ | $R = 7.5 \cdot 10^7 \cdot AHP^{1.6}$ | $R_m = 2.5 \cdot AHP^{0.9}$ | $C = \frac{9.7 \cdot 10^{-10}}{AHP^{0.6}}$ | $\tau = 6.2 \cdot 10^{-2} \cdot AHP$ | $I_{th} = \frac{1.8 \cdot 10^{-10}}{AHP^{1.7}}$ | | $ACV = \frac{2.7 \cdot 10^1}{AHP^{0.5}}$ |
| $ACV[m \cdot s^{-1}]$ | $S_{neuron} = 4.6 \cdot 10^{-10} \cdot ACV^{1.4}$ | $D_{soma} = 8.3 \cdot 10^{-8} \cdot ACV^{1.4}$ | $R = \frac{8.3 \cdot 10^{12}}{ACV^{3.5}}$ | $R_m = \frac{2.3 \cdot 10^3}{ACV^{2.1}}$ | $C = 9.0 \cdot 10^{-12} \cdot ACV^{1.4}$ | $\tau = \frac{7.4 \cdot 10^1}{ACV^{2.1}}$ | $I_{th} = 1.1 \cdot 10^{-15} \cdot ACV^{3.6}$ | $AHP = \frac{1.4 \cdot 10^3}{ACV^{2.2}}$ | |

#### Predicting correlations between MN properties

With this approach, the correlations between MN properties that were never concurrently measured in the literature are predicted, as displayed in Figure 2B. Such unknown relationships were indirectly extracted from the combination of known relationships (Table 3) and typical ranges of values obtained from the literature for these properties. Due to the prior five-fold cross-validation of the relationships in Table 4, these findings are reliable as indirectly consistent with the literature data processed in this study.

#### Extrapolation to rats and mice

It was assessed whether the scaled relationships obtained from cat data and reported in Table 4 could be successfully applied to rat and mouse data. A global {*I_th_; R*} dataset was obtained from 5 studies focusing on rats reported in Table TSM5, while four mice studies reported in Table TSM5 provided processable data for the pairs {*I_th_*; *R*}, {*τ; R*} and {C; *R*}. Only data from control groups of wild-type animals were considered while data on mutated, operated, or trained animals were disregarded. Because the information on the age distribution of the retrieved datasets is not available, data from adults rats and mice of unknown age were used, despite well-known age-related variations in the MN electrophysiological properties in these animals (Highlander et al., 2020; Huh et al., 2021). Like the cat data, low variability was observed between the *R* and *I_th_* distributions reported by different studies in the rat and mouse {*I_th_;R*} datasets. Figure 7 reports the global datasets for all 4 pairs of properties and the predictions obtained with the cat relationships reported in Table 4. As displayed in Table TSM5, 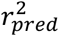 and 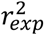 values were close for all 4 datasets, with a maximum 12.5% difference for {*τ;R*} in mice. *nRMSE* values remained low in both rat and mouse {*I_th_;R*} datasets, while being substantially higher in both {*τ;R*} and {*C;R*} datasets in mice because of an inter-species-specific offset in the relationships.

**Figure 7.**
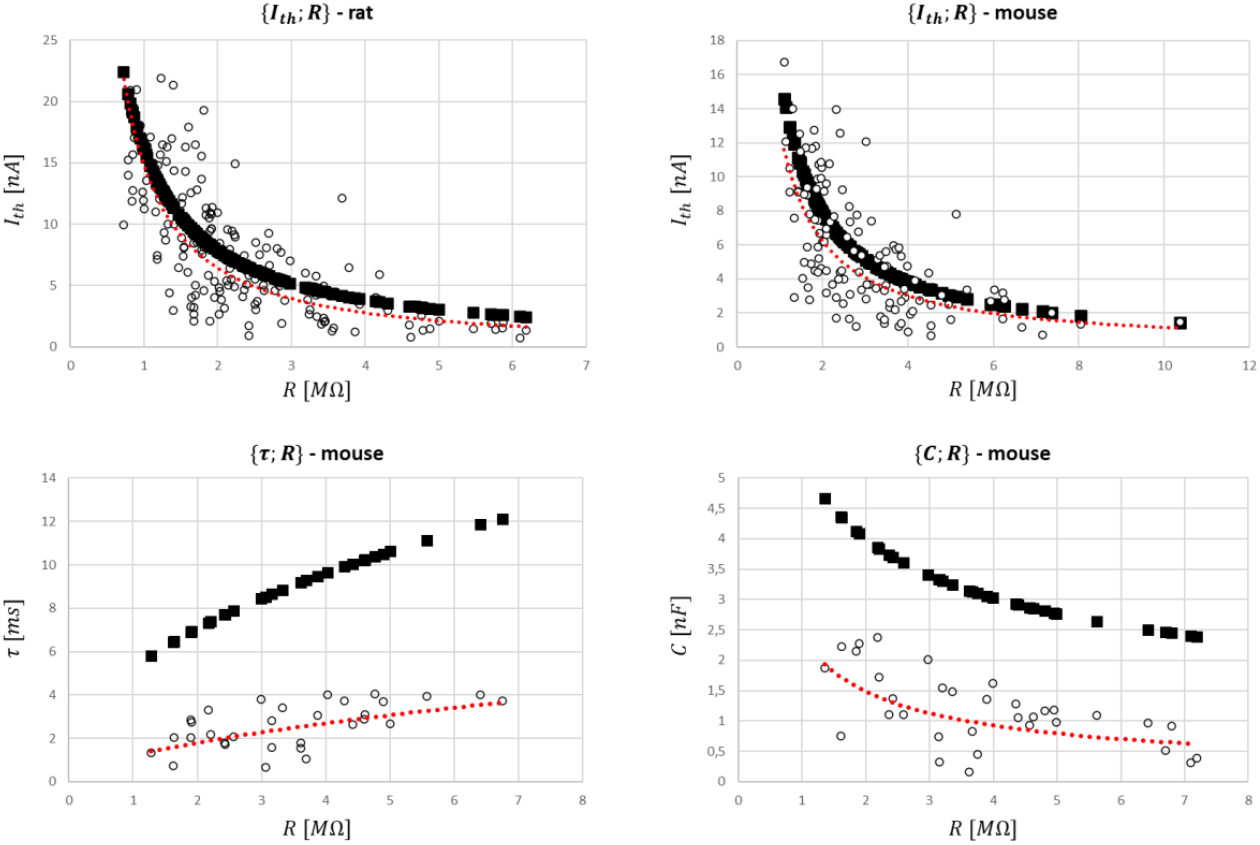
Global datasets for rat and mouse species and predictions of the MN properties with the cat mathematical relationships (Table 4). They were obtained from the 5 studies reporting data on rats and the 4 studies presenting data on mice reported in Table TSM5 that measured the {*I_th_*; *R*}, {*τ*; *R*} and {*C*; *R*} pairs of MN electrophysiological properties. *I_th_,R,τ,C* are given in *nA,MΩ,ms* and *nF* respectively. The experimental data (○ symbol) is fitted with a power trendline (red dotted curve) and compared to the predicted quantities obtained with the scaled cat relationships in Table 4 (■ symbol).

### Relationships between MN and mU properties

As shown in Figure 2A, eight pairs of one MN and one mU property were investigated in 14 studies in the literature in cats and rats. As remarked by Heckman & Enoka (2012) and Manuel et al. (2019), most recent studies were performed on mice, while most dated were dominantly performed on cats. One study on the rat gastrocnemius muscle (Kanda & Hashizume, 1992) indicated no correlation between *IR* and *CV*. However, after removing from the dataset 2 outliers that fell outside two standard deviations of the mean *IR* data, a statistically significant correlation (*p* < 0.05) between *IR* and *CV* was successfully fitted with a power trendline (*r*^2^ = 0.43). Eight studies, dominantly focusing on the cat soleus and medial gastrocnemius muscles, found a strong correlation between *F^tet^* and *CV*, while one study showed a significant correlation for the pair {*F^tet^;R*} in both the cat tibialis anterior and extensor digitorum longus muscles. One study (Burke *et al*., 1982) on the cat soleus, medial and lateral gastrocnemius muscles inferred a statistically significant correlation for {*F^tet^; S_MN_*}. In mice, both *R* and *I_th_* were significantly related to muscle tetanic and twitch forces. As *F^tet^, F^tw^* and *IR* are reliable indices of *S_mU_* as discussed in the Methods, the MN size-dependent relationships in Table 3 return eight 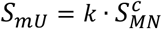 relationships between mU and MN properties (Table 5). A 2.8-fold range in positive c-values *c* ∈ [2.4; 6.6] (mean 3.8 ± 1.5 s.d.) was obtained between studies. This result infers than mU and MN size are positively related in cats, mice and potentially in rats.

**Table 5.**
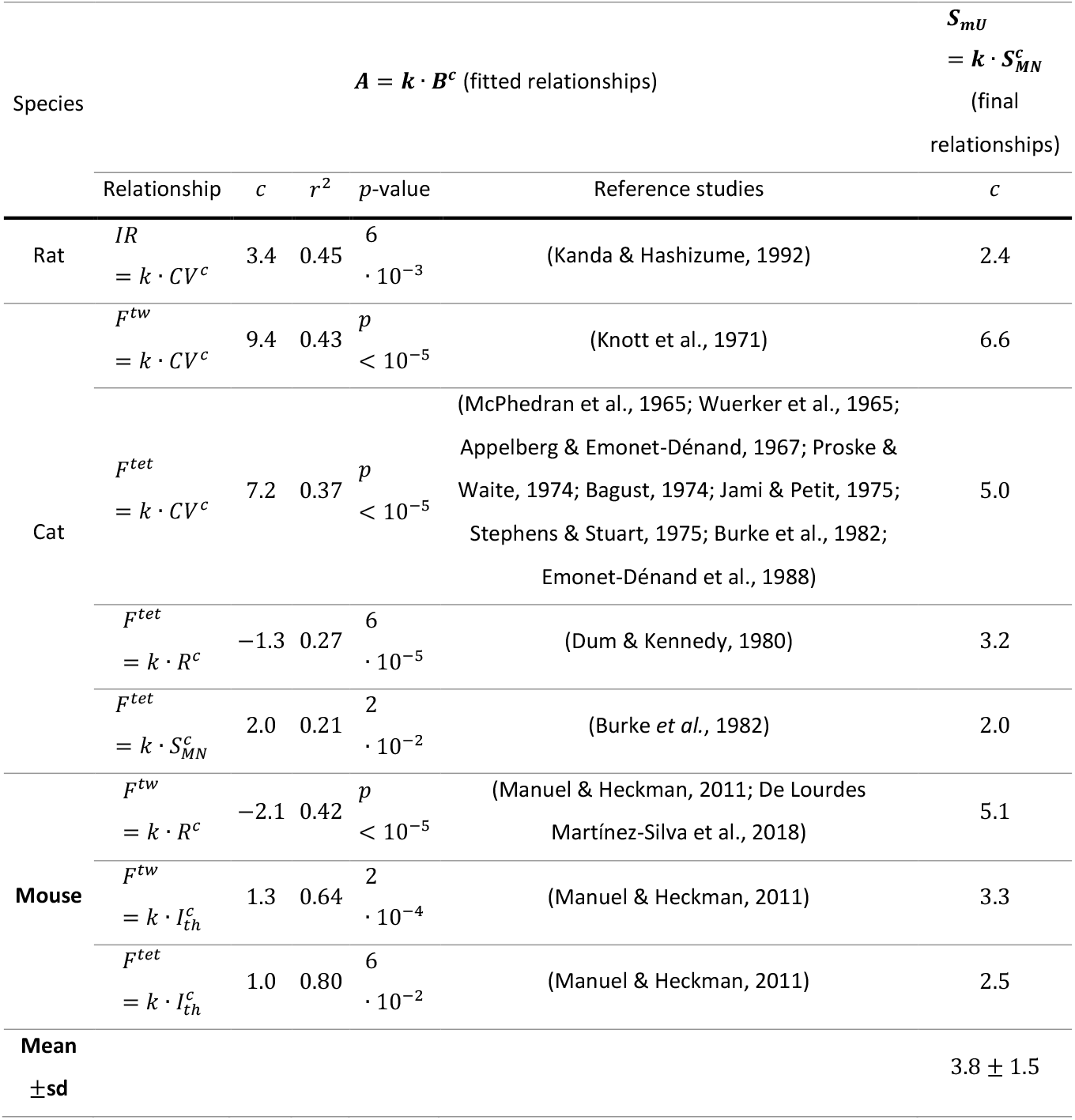
Fitted experimental data of pairs of one mU and one MN property and subsequent 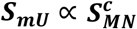 relationships. The *r*^2^, *p*-value and the equation *A* ∝ *B^c^* of the trendlines are reported for each pair of properties. In the last column, prior knowledge is used to derive 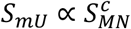 relationships.

## Discussion

We processed the normalized data (Figure 3) from previous cat experimental studies to extract mathematical relationships (Table 4) between several MN electrophysiological and morphological properties (Table 1), providing a clear summary of previously known qualitative inter-relations between these MN properties. Table 4 is a new convenient tool for neuroscientists, experimenters, and modelers. Besides obtaining significant quantitative correlations between MN electrophysiological properties and MN size (Table 3, Figure 4, Figure 5), we established (Table 4) and validated (Figure 6) a comprehensive mathematical framework that models an extensive set of experimental datasets available in the literature, quantitatively links all the pairs of the MN properties in Table 1, and is consistent with Rall’s cable theory, as discussed in the following. This framework is directly applicable, although with limitations, to other mammalian species (Figure 7), and is used along with other measurements of mU properties (Table 5) to discuss the Henneman’s size principle.

### Consistency of the relationships with previous empirical results and Rall’s theory

The mathematical relationships derived in Table 4 are first in strong agreement with the relative variations between MN properties that were reported in review papers and are listed in the Introduction section. Some combinations of the relationships in Table 4 are besides consistent with other cat experimental data that were not included in the global datasets and were not used for deriving the relationships. For example, the relationships 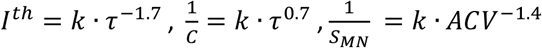 and 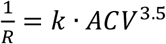 in Table 4 yield, when combined, 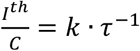 and 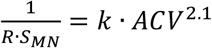, which are consistent with the relationship 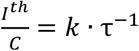 experimentally observed by Gustafsson & Pinter (1985) and 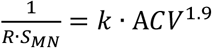 measured by Kernell & Zwaagstra (1981). The stronger-than-linear inverse relationship 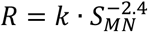 is also consistent with the phenomenological conclusions by Kernell & Zwaagstra (1981). Moreover, some standard equations resulting from Rall’s cable theory can be derived with a combination of the relationships in Table 4, assuming the MNs as equivalent cylinders with spatial uniformity of both *C_m_* and *R_m_*. For example, taking 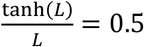, which is in the typical [0.48; 0.76] range reported in the literature (Powers & Binder, 2001), the relation 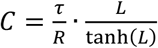 (eq.1. in Gustafsson & Pinter (1984a)) is obtained when combining the *R* – *S_neuron_, C* – *S_neuron_* and *τ* – *S_neuron_* relationships. Also, taking 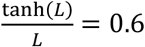 (standard value provided in Powers & Binder (2001)), the *R* – *S_neuron_* equation in Table 4 applied to 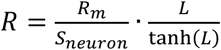 in Rall (1977) yields 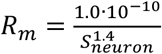. This *R_m_* – *S_neuron_* relationship is consistent with the observations in Powers & Binder (2001) that smaller MNs have larger *R_m_* values. Also, considering that 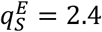 and *S_neuron_* ∈ [0.18; 0.44] *mm*^2^ (Table TSM3), 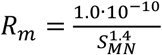 makes *R_m_* vary over the 3.5-fold theoretical range [0.12; 0.44] Ω · *m*^2^ in the MN pool. This is highly consistent both with the 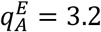-fold range and the 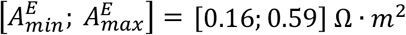 mean range previously reported in the literature (Albuquerque & Thesleff, 1968; Barrett & Crill, 1974; Burke et al., 1982; Gustafsson & Pinter, 1984b). Also, 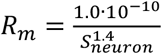 yields *R_m_* = 2.5 · *AHP*^0.9^ from the relationships in Table 4, which is consistent with the indirect conclusion of a positive correlation between *R_m_* and *AHP* in Gustafsson & Pinter (1984b). Finally, 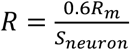 and 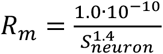 support the statement that the variations in *R_m_* in the MN pool may be as important as cell size *S_neuron_* in determining MN excitability (Gustafsson & Pinter, 1985), although the variation of *R* in a MN pool may be also related to cell geometry (Gustafsson & Pinter, 1984b), which was not investigated in this study. This is consistent with the results in Table TSM4 in Supplementary Material, which demonstrate that the range of variation 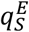 of *S_MN_* does not entirely explain the ranges of variation 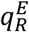 and 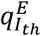 of *R* and *I_th_*, and that MN excitability is not only determined by MN size. Because of the aforementioned consistency with previous findings and with Rall’s theory, the remaining relationships between *R_m_* and the other MN properties were calculated from 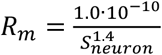, following the same Methods as before, and added to Table 4. Importantly, *R_m_* might however not be uniform across the somatodendritic membrane, according to results from completely reconstructed MNs (Fleshman et al., 1988), and might be positively correlated to the somatofugal distance (Fleshman et al., 1983), with dendritic *R_m_* being higher than somatic *R_m_* (Barrett & Crill, 1974; Powers & Binder, 2001; Kernell, 2006) by a factor 100-300 (Clements & Redman, 1989). This contradicts the assumptions of membrane isopotentiality necessary to apply Rall’s cable theory. The relationships in Table 4, which were obtained from measurements in the soma where *R_m_* is mainly constant, may thus not directly extrapolate to the dendritic regions of the MNs. Yet, the dimensionless variations of the average across MN surface of *R_m_* with the other MN properties may still be valid (Kernell, 2006).

The equations in Table 4 support the notion that MNs behave like resistive-capacitive systems, as demonstrated by the combination of the *R* – *τ* and *C* – *τ* relationships in Table 4, which yields *τ*^0.9^ = 0.8 · *RC*, close to the theoretical equation 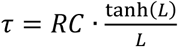 derived from Rall’s theory. Also, Table 4 directly yields, from the combination of 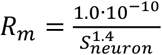 with the other relationships, *τ* = 1.4 · 10^−2^ · *R_m_*, which is exactly eq.2.15 in Rall (1977) (*τ* = *R_m_C_m_*) with a constant value for *C_m_* = 1.4 · 10^−2^*F* · *m*^−2^ that is consistent with the literature.

The relationship *C* = 1.3 · 10^−2^ · *S_neuron_* obtained in Figure 4 and reported in Table 4 is in agreement with the widely accepted proportional relationship between MN membrane capacitance and surface area, when spatial uniform values of *R_m_* and *C_m_* are assumed. It is worth noting that this proportional relationship was obtained in this study without simultaneous empirical measurements of *S_MN_* and *C*. The values of *C* were obtained from Rall’s 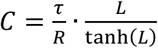 equation (Gustafsson & Pinter, 1984b) and the corresponding values of *S_MN_* were indirectly obtained from the final datasets {*ACV*; *S_MN_*}, {*AHP; S_MN_*}, {*R; S_MN_*} and {*I_th_*;*S_MN_*} that involved the data from 17 selected studies as described in Figure 1. Although consistent with typical values reported in the literature (Lux & Pollen, 1966; Albuquerque & Thesleff, 1968; Barrett & Crill, 1974; Adrian & Hodgkin, 1975; Sukhorukov et al., 1993; Major et al., 1994; Solsona et al., 1998; Thurbon et al., 1998; Gentet et al., 2000), the *C_m_* = 1.3 · 10^−2^*F* · *m*^−2^ constant value reported in this study is slightly higher than the widely-accepted *C_m_* = 1.0 · 10^−2^*F* · *m*^−2^ value (Ulfhake & Kellerth, 1984; Gustafsson & Pinter, 1984b). This high value is consistent with the conclusions from Gustafsson & Pinter (1984a) who obtained slightly conservative values for *S_neuron_* when approximating the MN surface area as 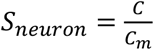 with *C_m_* = 1.0 ·10^−2^*F* · *m*^−2^.

Finally, the empirical relationship 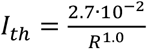 (Table 4) provides Δ*V_th_* = 27*mV*, consistently with typical reported values (Brock et al., 1952; Eccles et al., 1958), despite uncertainties in the value of the membrane resting potential (Heckman & Enoka, 2012). This supports the conclusions that the relative voltage threshold Δ*V_th_* may be assumed constant within the MN pool in first approximation (Coombs et al., 1955; Gustafsson & Pinter, 1984a; Powers & Binder, 2001), and that Ohm’s law is followed in MNs for weak subthreshold excitations (Glenn & Dement, 1981; Ulfhake & Kellerth, 1984; Spruston & Johnston, 1992; Powers & Binder, 2001; Kernell, 2006), as discussed in the Methods Section. However, voltage thresholds Δ*V_th_* tend to be lower in in MNs of large *R* and long *AHP* (Gustafsson & Pinter, 1984a), suggesting that Δ*V_th_* might be inversely correlated to MN size *S_MN_*. Moreover, because of the voltage-activation of persistent inward currents near threshold which add to the external stimulating current and increases the MN input resistance (Fleshman et al., 1988), with a prominent effect on small MNs (Powers & Binder, 2001), the experimental values of Δ*V_th_* are generally greater than that directly predicted from the product *I_th_* · *R* (Gustafsson & Pinter, 1984b), and a variant of Ohm’s law such as ΔV_th_(*V*) = *I_th_* · *R*(*V*) should be considered during MN discharging events. This voltage-dependent variation in the values of *R* might partly explain why the range of the values of *R* reported in the literature for weak subthreshold currents (Table TSM4) is generally lower than that of *I_th_*. While ΔV_th_ may be always size-dependent, and size-voltage-dependent close to threshold and during firing events, no correlation between 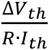 and *C, AHP* or *ACV* was found and reported in Table 4 in subthreshold conditions, consistent with measurements performed by Gustafsson & Pinter (1984b) and Ulfhake & Kellerth (1984), substantiating that the dynamics of MN recruitment dominantly rely on *R,I^th^* and Δ*V_th_* (Heckman & Enoka, 2012).

### Relevance of the relationships

Table 4 provides the first quantitative description of an extensive set of cat experimental data available in the literature. These inter-related relationships were validated, successfully reconciliate the conclusions formulated by the selected experimental studies and are in strong agreement with the theoretical equations describing Rall’s cable theory. This robust framework advances our general understanding of the MN neurophysiology by inferring quantitative relationships for pairs of MN properties that were never concurrently measured in experiments (Figure 2B) by crossmatching other relationships involving these properties. For example, 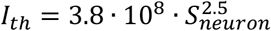 (Table 4) quantitatively supports the size-dependency of *I_th_* that was predicted by Powers & Binder (2001) from Rall’s equations. Furthermore, modelers can directly and realistically tune with Table 4 the physiological parameters of phenomenological MN models, such as leaky fire-and-integrate models, and/or build pools of MNs displaying a continuous distribution of realistic MN profiles of MN-specific electrophysiological and morphometric properties, as it has been previously attempted (Dong et al., 2011; Negro et al., 2016). Models tuned with Table 4 can for example better replicate the discharging behaviour of MNs, obtained from decomposed high-density EMGs, than models scaled with generic property values (Caillet et al., 2022). In this view, such tuned models display MN-specific behaviours that should be easier to interpret. The mathematical relationships in Table 4 can also support future experimental investigations performed on spinal MNs in adult mammals *in vivo* by completing experimental datasets for which properties that are difficult to measure were not obtained, such as the MN membrane time constant. Experimenters can therefore choose to reduce the workload of measuring all the MN properties in Table 1 to favour the identification of a maximum number of MNs and obtain extensive datasets describing larger MN populations that provide an informative window on the continuous distribution of MN properties in a MN pool. The unknown electrophysiological MN properties, typically obtained *in vivo*, of cadaveric specimens can also be estimated with the size-related relationships in Table 4 from in vitro measurements of the somal diameter *D_soma_* in a slice preparation of the spinal cord. The mathematical relationships in Table 4 describe the experimental data published in the literature and provide a convenient metric for experimenters to compare their measurements to previous findings.

### Extrapolation of the relationships to other mammals

It is demonstrated in Figure 7 and with the calculation of the *r*^2^ values in Table TSM5 that the mathematical equations in Table 4 obtained from cat data adequately predict the normalized associations between pairs of MN properties in rats and mice. However, the scaling factor *K* applied to the intercept *k_e_* to scale the normalized relationships *A* = *K* · *k_e_* · *B^e^* is species-specific, being for example around 3 times smaller in mice than in cat for the {*τ;R*} and {*C;R*} pairs (Figure 7) and explaining the large *nRMSE* values in these cases. Absolute values of the {*I_th_*; *R*} pair were nevertheless accurately predicted in both rats and mice from the cat relationships, as explained by the larger values of *R* which counterbalances the respectively lower *I_th_* values in mice and rats than in cats (Manuel et al., 2019; Highlander et al., 2020). Despite the age-related data variability (Highlander et al., 2020) in the rodent dataset displayed in Figure 7, these findings advance our understanding of the systematic inter-mammalian-species variations in MN properties. While the mathematical equations in Table 4 are specific to cats, the normalized equations in Supplementary Material can be scaled with species-specific data if investigating other mammals.

### Henneman’s size principle of motor unit recruitment

Table 5 reports statistically significant power relationships of positive power values between MN and mU indices of size. These results substantiate the concept that *S_MN_* and *S_mU_* are positively correlated in a motor unit (MU) pool and that large MNs innervate large mUs (Henneman, 1981; Heckman & Enoka, 2012), a statement that has never been demonstrated from the concurrent direct measurement of *S_MN_* and *S_mU_*. Besides, considering the positive *I_th_ – S_MN_* correlation (Table 4), and that the mU force recruitment threshold *F^th^* is positively correlated to *F^tet^* (Heckman & Enoka, 2012) and thus to *S_mU_* (Table 2), larger MUs have both larger current and force recruitment thresholds *I_th_* and *F^th^* than relatively smaller MUs, which are thus recruited first, consistently with the Henneman’s size principle of MU recruitment (Henneman, 1957; Wuerker et al., 1965; Henneman et al., 1965a; Henneman et al., 1965b; Henneman et al., 1974; Henneman, 1981; Henneman, 1985). The terminologies ‘small MU’, ‘low-force MU’ and ‘low-threshold MU’ are thus equivalent. Henneman’s size principle thus relies on the amplitude of the MN membrane resistance (Binder et al., 1996; Powers & Binder, 2001; Heckman & Enoka, 2012). Finally, the relationships 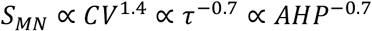 (Table 4) suggest that high-threshold MUs rely on relatively faster MN dynamics, which might partially explain why large MNs can attain relatively larger firing rates than low-thresholds MNs, for example during ballistic contractions or during events close to maximum voluntary contractions (Powers & Binder, 2001; Heckman & Enoka, 2012).

It has been repeatedly attempted to extend Henneman’s size principle and the correlations between the MU properties in Table 1 to the concept of ‘MU type’ (Burke & Ten Bruggencate, 1971; Burke, 1981; Bakels & Kernell, 1993; Powers & Binder, 2001). While a significant association between ‘MU type’ and indices of MU size has been observed in some animal (Fleshman et al., 1981; Burke et al., 1982; Zengel et al., 1985) and a few human (Milner-Brown et al., 1973; Stephens & Usherwood, 1977; Garnett et al., 1979; Andreassen & Arendt-Nielsen, 1987) studies, it has however not been observed in other animal studies ((Bigland-Ritchie et al., 1998) for a review) and in the majority of human investigations (Sica & McComas, 1971; Goldberg & Derfler, 1977; Yemm, 1977; Young & Mayer, 1982; Thomas et al., 1990; Nordstrom & Miles, 1990; Elek et al., 1992; Macefield et al., 1996; Van Cutsem et al., 1997; Mateika et al., 1998; Fuglevand et al., 1999; Keen & Fuglevand, 2004). Moreover, the reliability of these results is weakened by the strong limitations of the typical MU type identification protocols. Sag experiments are irrelevant in humans (Buchthal & Schmalbruch, 1970; Thomas et al., 1991; Bakels & Kernell, 1993; Macefield et al., 1996; Bigland-Ritchie et al., 1998; Fuglevand et al., 1999), and lack consistency with other identification methods (Nordstrom & Miles, 1990). MU type identification by twitch contraction time measurements is limited by the strong sources of inaccuracy involved in the transcutaneous stimulation, intramuscular microstimulation, intraneural stimulation, and spike-triggered averaging techniques (Taylor et al., 2002; Keen & Fuglevand, 2004; McNulty & Macefield, 2005; Negro et al., 2014; Dideriksen & Negro, 2018). Finally, as muscle fibres show a continuous distribution of contractile properties among the MU pool, some MUs fail to be categorized in discrete MU types in some animal studies by histochemical approaches (Reinking et al., 1975; Totosy de Zepetnek et al., 1992). Owing to these conflicting results and technical limitations, MU type may not be related to MN size and the basis for MU recruitment during voluntary contractions (McNulty & Macefield, 2005; Duchateau & Enoka, 2011).

### Limitations

The mathematical relationships derived Table 4 and the conclusions drawn in the Discussion present some limitations, most of which were previously discussed and are summarised here. A first limitation relates to the experimental inaccuracies in measuring the MN properties in Table 1. As discussed, the true values of *R* and *τ*, and thus of *R_m_* and *C*, may be underestimated because of a membrane leak conductance around the electrode and some voltage-activated membrane nonlinearities. Yet, the selected studies reduced the impact of these phenomena, which besides may not affect the association between MN properties (Gustafsson & Pinter, 1984a). Also, *τ, C* and *R_m_* were estimated assuming that the MN membrane is isopotential. This is valid in the soma, where *τ, C* and *R_m_* were measured, and the relationships in Table 4 are adequate for a MN simplified to one equivalent cable. However, as the membrane resistivity may increase with somatofugal distance, as discussed previously, the relationships in Table 4 may be offset towards larger *R* values in dendritic areas, which should be accounted for when modelling the MN dendrites in compartmental MN models or completely reconstructed dendritic trees. Finally, the selected cat studies are relatively dated, as observed in Manuel et al. (2019). Thus, recent computer-assisted techniques of MN morphometric measurements were notably not applicable in those studies, yielding important sources of inaccuracies, as discussed in Supplementary Material SM2. Therefore, the relationships involving *S_neuron_* are subjected to a higher level of inaccuracy. A second limitation arises from the inter-study variability of the experimental datasets reported in the selected studies. As discussed, animal preparations slightly diverged between the selected studies. Also, because of the scarcity of available data in the literature, measurements obtained from MNs innervating different muscles from adult animals of different sex and age were merged into unique datasets. This inter-study variability, added to the ‘unexplained’ variance reported in Highlander et al. (2020), was however reduced by the normalization of the experimental datasets, as investigated in Figures FSM2 and FSM4 in Supplementary Material. The normalization approach is however only valid if the experimental studies identified the same largest MN relatively to the MN populations under investigation. This cannot be verified but is likely as most studies returned similar normalized distributions of their measured parameters, as displayed in Figure FSM1 (Supplementary Material), inferring that a similar portion of the MN pools was identified in the selected studies. Finally, the results proposed in this study may be affected by methodological bias from the three research groups that provided most of the data points processed in this study, even if all the selected studies reported similar methods to measure the properties, as discussed in the Methods section.

A third limitation is due to the limited processable experimental data available in the literature, especially in rats and mice, as also discussed by Heckman & Enoka (2012). The extrapolation of the relationships in Table 4 to rat and mouse species could only be assessed against a small set of studies and for three associations only. The conclusions on the associations between *S_mU_* and *S_MN_* also relied on few studies providing few measurements, due to the considerable amount of work required to measure both MN and mU properties in single MUs. In cats, some pairs of MN properties were investigated in only one study, preventing inter-study comparisons, and stressing the usefulness of Table 4 for reconciliating the data available in the literature. Within the experimental datasets, the scarcity of data points in some regions of the data distributions (see the skewed *I_th_* and *R* distributions in the density histograms in Figure FSM1 for example), transfers during processing to the final size-dependent datasets (Figure 4). This data heterogeneity may affect the reliability of the relationships in Table 4 for the extreme MN sizes (see, in Figure 4, the 95% confidence interval of the regressions widening for small *S_MN_* in the {*AHP; S_MN_*} and {*R; S_MN_*} datasets and for large *S_MN_* in the {*I_th_; S_MN_*} dataset). In the validation procedure, the highest *nME* values reported in Figure 6 were obtained for these regions of limited data for most of the global datasets. This heterogeneous density of the data may be explained by the natural skewness of the MN property distributions in the MN pool towards many small MNs and few large MNs (Heckman & Enoka, 2012) and/or a systematic experimental bias towards identifying and investigating relatively large MUs.

A fourth limitation arises from the relatively low *r*^2^ values obtained from the power regressions in Table 3. Although these power trendlines are statistically significant (see p-values in Table 3), and globally provide a better description of the data than other fits (see Table TSM1), they cannot entirely describe the associations between MN property distributions, in line with the conclusions reported in the selected studies. According to these first four limitations, the mathematical relationships in Table 4 best reproduce published cat data but include the level of inaccuracies of the original experimental approaches.

A fifth limitation is the restriction of the study to the MN properties reported in Table 1. Other MN-specific properties, such as the electronic length, the relationship between discharge rate and excitatory current and its hysteresis, or the amplitude of the inhibitory and excitatory postsynaptic potentials were omitted because they were rarely measured or reported along with *S_MN_,ACV,AHP,R,I_th_,C* and *τ* in the selected studies. Moreover, as the cell geometry was not investigated in this study, other MN properties (Clements & Redman, 1989) such as the effects of the distributed synaptic integration along the dendritic tree were overlooked. Finally, apart from *I_th_,ACV* and *AHP*, all the MN properties investigated in this study are passive properties, i.e. base-line properties of the cell at rest (Powers & Binder, 2001), as they were measured using weak sub-threshold current pulses. Therefore, other MN active properties, such as voltage-dependent ionic channel-related properties including persistent inward currents (PICs), which activate close or above threshold, were not considered in this study. While this limits the applicability of Table 4 to a functional context or in comprehensive Hodgkin-Huxley-like models of MNs, some relationships in Table 4 remain pertinent in those conditions if they are linked with other observations in the literature on MN active properties, e.g., MNs of small *S_MN_* having longer lasting total PICs of greater hyperpolarized activation than MNs of large *S_MN_* (Heckman & Enoka, 2012). Despite the aforementioned limitations, the relationships in Table 4 can be directly used in phenomenological RC approaches like LIF models (Izhikevich, 2004; Teeter et al., 2018), which rely on the properties reported in Table 1, to derive profiles of inter-consistent MN-specific properties and to describe realistic continuous distributions of the MN properties *R, I_th_, C, ΔV_th_* and *τ* in the MN pool, as attempted in previous works (Dong et al., 2011; Negro et al., 2016). An example of a computational modelling application of Table 4 is provided in Caillet et al. (2022), where the relationships tune a cohort of LIF models which estimate, from the firing activity of a portion of the MN pool obtained from decomposed high-density EMGs, a realistic distribution of the MN properties in the MN pool and the firing behaviour of a complete MN pool in an isometrically contracting human muscle.

As a final limitation, the relationships in Table 4 were obtained from a regression analysis and therefore provide correlations between some MN properties in a MN pool but cannot be used to draw conclusions on the causality behind these associations.

### Conclusion

This study provides in Table 4 a mathematical framework of quantitative associations between the MN properties *S_MN_, ACV, AHP, R, R_m_, I_th_, C* and *τ*. This framework, which is consistent with most of the empirical and theoretical conclusions in the literature, clarifies our understanding of the association between these MN properties and constitutes a convenient tool for neuroscientists, experimenters, and modelers to generate hypothesis for experimental studies aiming at investigating currently unreported relationships, support experimentations and build virtual MN profiles of inter-consistent MN-specific properties for MN modelling purposes.

## Supporting information

data for Figure 7

data for Table 5

data for Figure 3

data for Figure 7

## ACKNOWLEDGEMENTS

AC was supported by the Skempton Scholarship and LM by an Imperial College Research Fellowship granted by Imperial College London. The authors want to thank Dr Marin Manuel for making available the experimental data published in Huh et al., 2021, Time course of alterations in adult spinal motoneuron properties in the SOD1 (G93A) mouse model of ALS, Eneuro.

