## supplementary materials for "Mathematical Relationships between Spinal Motoneuron Properties"

### SM1 – Inter-study data variability

Inter-study variability greater than the threshold defined in the Methods section was observed in 3,2 and 4 property distributions for the metrics Range, Mean and CoV metrics, respectively. Indeed, sensibly different ranges of property values were observed to be reported between studies of the $\{ACV; S_{MN}\}$, $\{AHP; ACV\}$, and $\{R; AHP\}$ datasets for the $S_{MN}$, $ACV$, and $AHP$ properties, respectively, with $sd_g \in [10; 15]\%$ and/or $\frac{sd_g}{mean_g} \in [0.15; 0.33]$. Also, the studies of the $\{R; AHP\}$ and $\{\tau; R\}$ datasets returned some variability in the mean of the reported data for properties $AHP$ and $R$, respectively, with $sd_g \in [10; 11]\%$ and/or $\frac{sd_g}{mean_g} \in [0.15; 0.23]$. The $\{ACV; S_{MN}\}, \{R; ACV\}$ and $\{R; AHP\}$ global datasets showed variable data dispersion around the mean for properties $S_{MN}$ , $R$ and $ACV$, and $AHP$, respectively, with $sd_g \in [10; 11]\%$ and/or $\frac{sd_g}{mean_g} \in [0.15; 0.30]$.

### SM2 – Discussion on the potential underestimation of $q_S^E$

The minimum and maximum values of $S_{neuron}$ and $D_{soma}$ reported in Table TSM1 may underestimate the true distribution of MN sizes in a MN pool for several reasons, and thus underestimate the true value of $q_S^E$. First, the studies reported in Table TSM1 investigated MN populations of small size, so that the largest and smallest MNs of the investigated MN pools may not have been systematically identified and measured. Also, dendritic labelling by staining may only identify dendrites up to the 3rd dendritic branch (Issa et al., 2010; Fogarty et al., 2018) and full dendritic trees may not be identified, thus underestimating $S_{neuron}$ for the largest MNs that exhibit the highest dendritic tree complexity (Fogarty et al., 2020). Finally, the distribution of gamma- and alpha-MN sizes is typically bimodal and the size distributions of the two populations are reported to overlap (Moschovakis et al., 1991; von Steyern et al., 1999; Friese et al., 2009; Deardorff et al., 2013). However, most studies reported in Table TSM1 identify the alpha MNs from the gamma MNs by visually splitting the bimodal distributions of MN sizes in two independent domains (Donselaar et al., 1986; Ishihara et al., 2001), thus neglecting the overlap. Similarly, some studies identify as gamma MNs any MN showing an axonal conduction velocity typically less than $\sim 60 m \cdot s^{-1}$ (Zwaagstra, B. & Kernell, 1980; Ulfhake, B. & Kellerth, 1984). These approaches lead to overestimating the lower limit for the reported $D_{soma}$ and $S_{neuron}$ values. Consequently, true ranges in MN sizes may be larger than the theoretical ones reported in Table TSM1. A similar comment can be made for the electrophysiological properties reported in Table TSM2. However, the $q_S^E = 2.4$-fold range obtained for cats is consistent with the fold-ranges obtained from rat and mouse data for $D_{soma}$, as reported in Table TSM1, while it underestimates the 4.4-fold range in $S_{neuron}$ values observed in mice. Table TSM1 finally shows that typical cat MN sizes are larger than in rats and mice, as reviewed by Manuel et al. (2019).

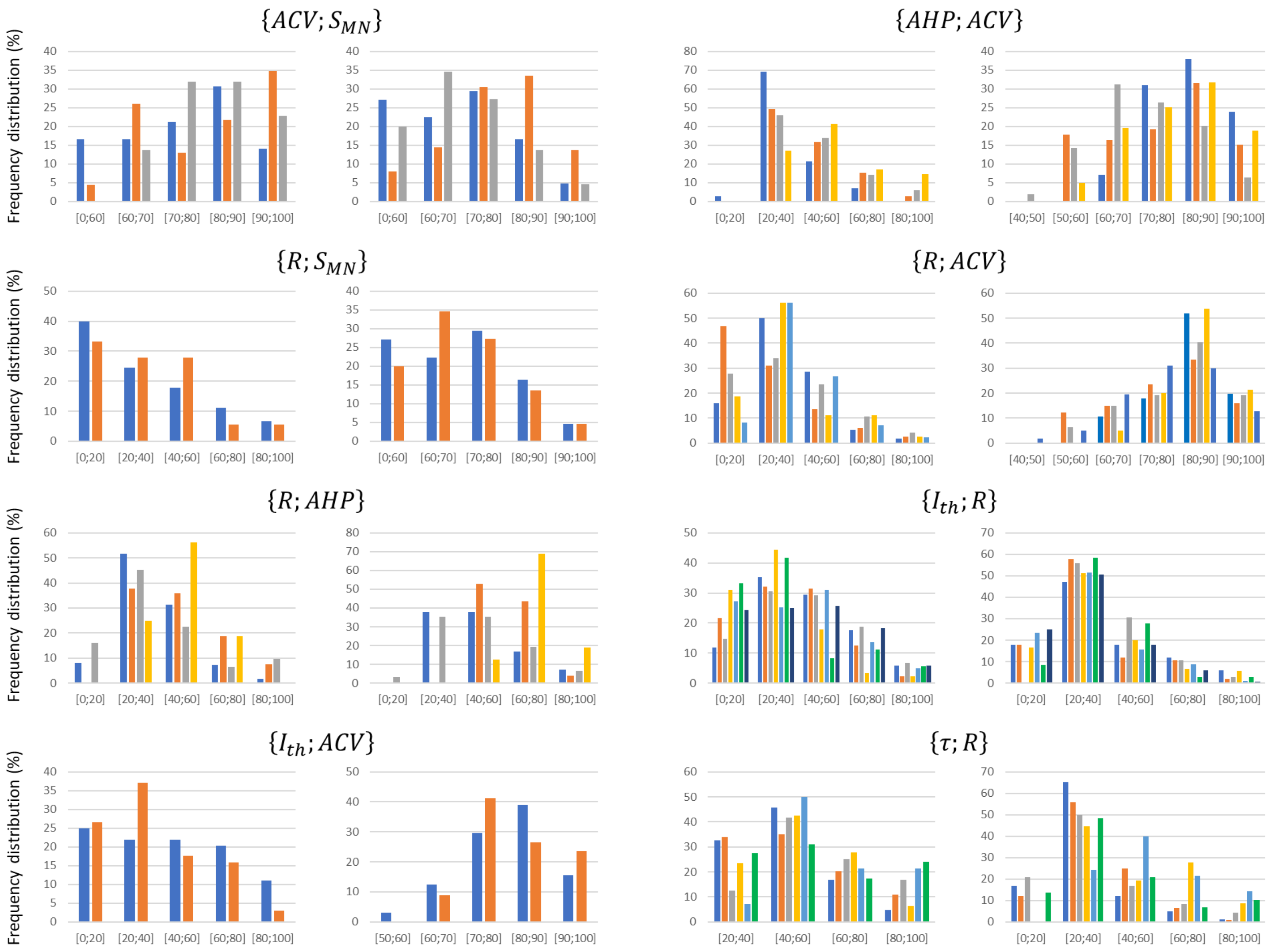


**Figure FSM1. Density histograms of the data distributions reported in the experimental studies included in the global datasets.** Depending on the typical range over which each property spans, the distributions are divided in steps of 10% or 20%. The frequency distribution is provided in percentage of the total number of reported data points in a study. Different studies are displayed with different colours in each graph. For each dataset $\{A; B\}$, the frequency distributions of property $A$ and $B$ are provided in the left and right plot respectively.

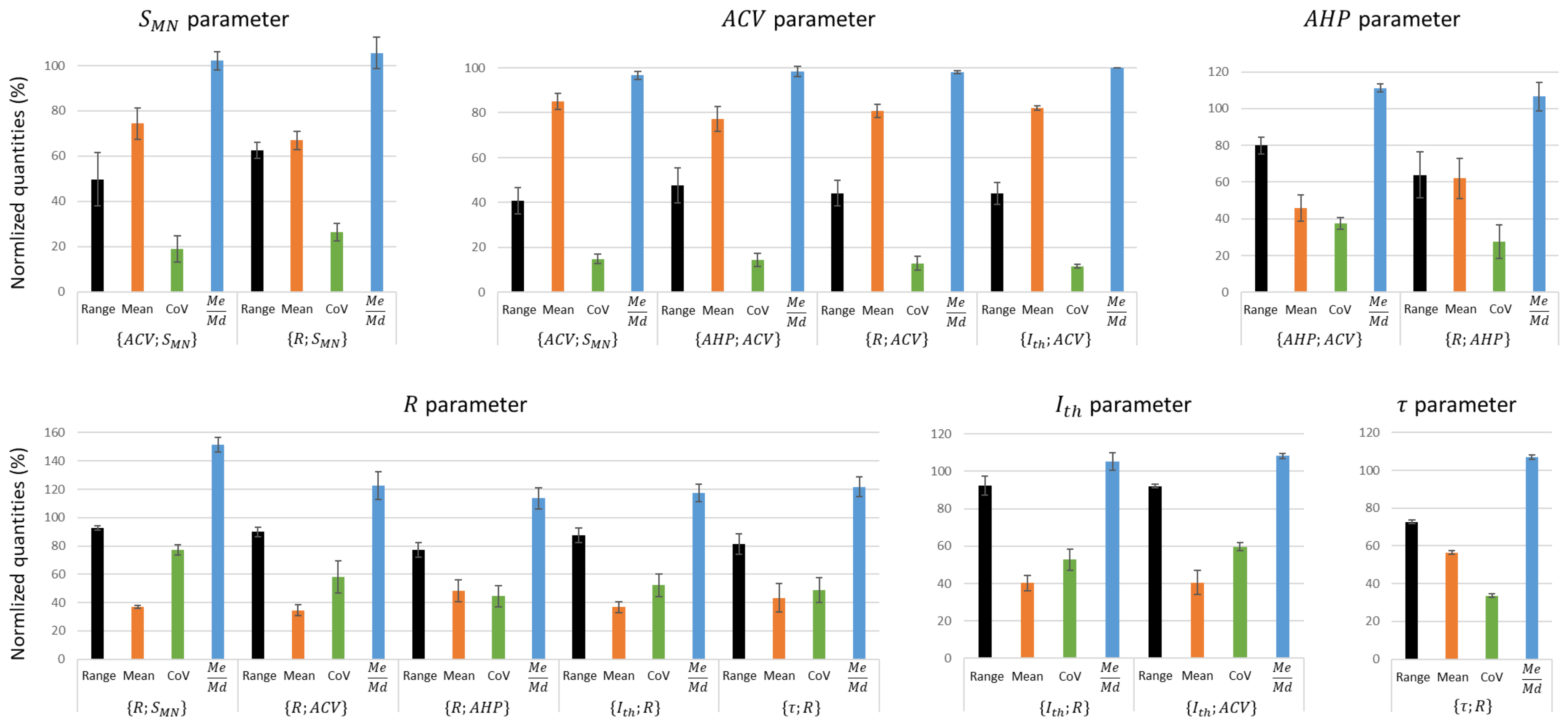


**Figure FSM2. Assessment of data variability between the experimental studies that constitute the 8 global datasets that include at least two experimental studies.** For each experimental study included in the global dataset $\{A; B\}$, the Range, Mean, coefficient of variation ($CoV = \frac{standard\ deviation}{Mean}$) and the ratio $\frac{Me}{Md} = \frac{Mean}{Median}$ of the experimental $A$ values measured in this study were computed. Then, the average (bars) and standard deviation (error bars) across experimental studies of these four metrics (Range, Mean, CoV, $\frac{Me}{Md}$) were calculated for each global dataset independently. For example, the first bar in the '$S_{MN}$ parameter' plot represents the average across three studies (Kernell & Zwaagstra, 1981; Cullheim, 1978; Burke et al., 1982) of the range of $S_{MN}$ values reported in these studies, for the global dataset $\{ACV; S_{MN}\}$. The computed standard deviations (error bars) express for each global dataset the inter-study variability of (1) the length of the identified bandwidth of the MN pool (Range metric), (2) the spread of values around the mean (CoV metric), (3) the skewness of the distributions, and (4) whether the distributions from different studies are centred (Mean metric). A global dataset $\{A; B\}$ reporting narrow error bars for parameter $A$ for the four metrics infers that the experimental studies constituting this dataset measured similar distributions of property $A$.

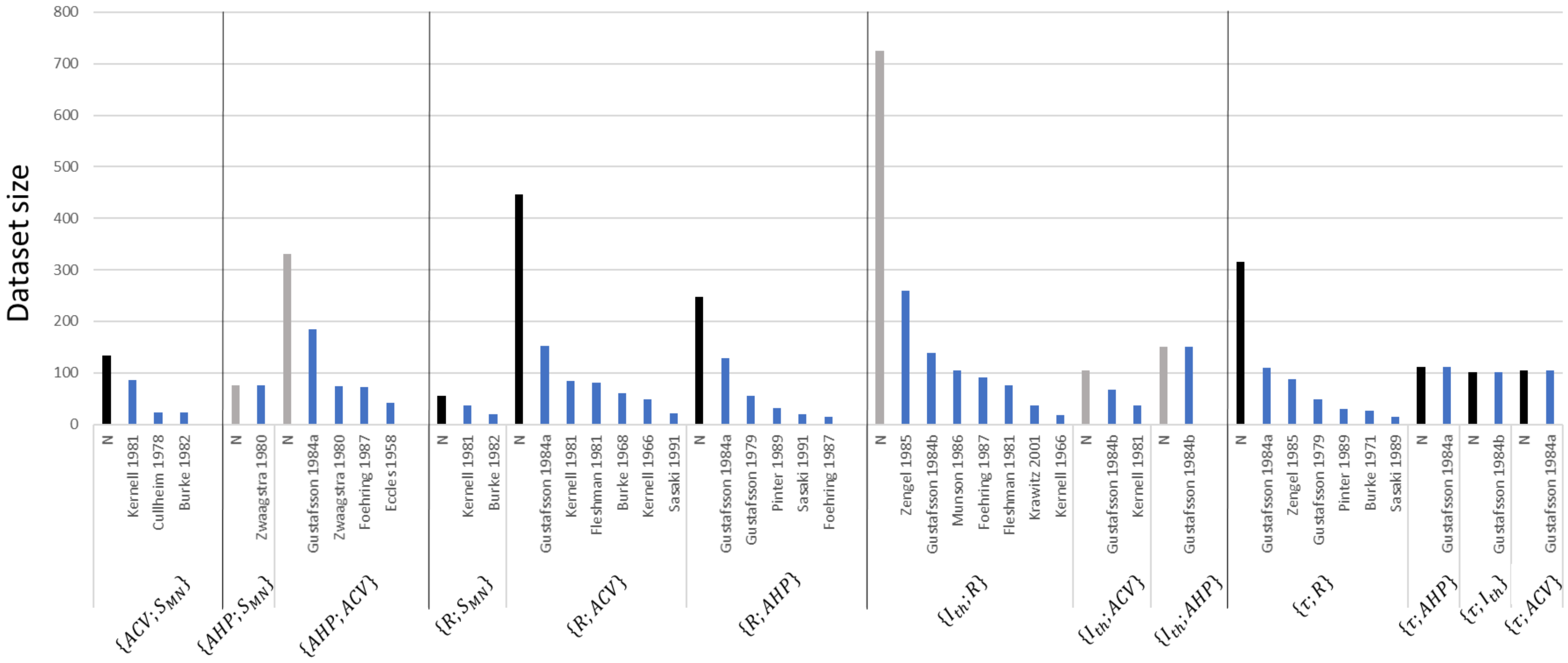


**Figure FSM3. Distribution of the size of the experimental datasets constituting the global datasets for assessment of the variability in the input data.** The histogram is divided between global datasets (half vertical lines), grouped as final size-dependent datasets (full vertical lines). For each global dataset, the total number of data points is reported (label: "N"), while the size of the constitutive experimental studies is reported in decreasing order.

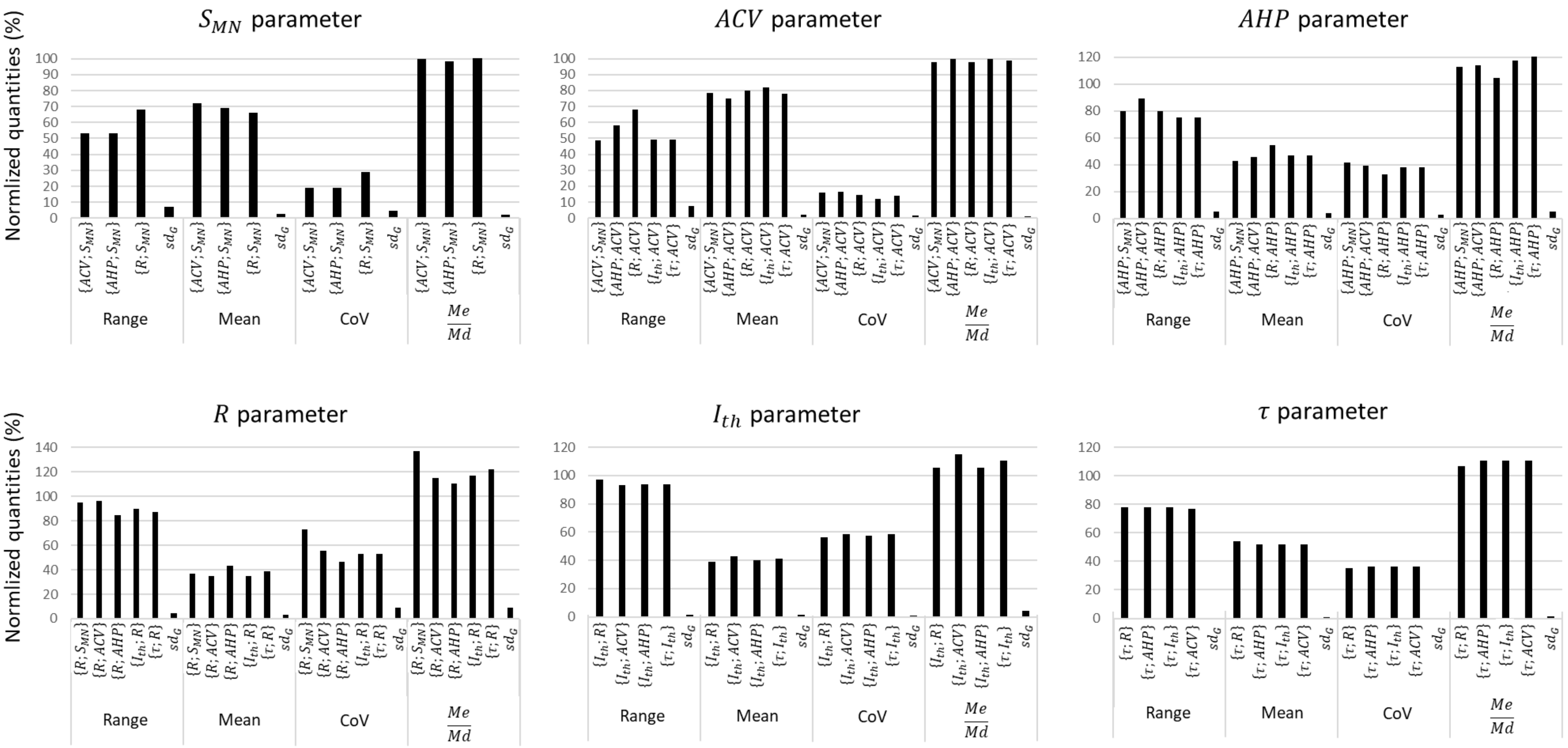


**Figure FSM4. Assessment of data variability between the global datasets.** This figure compares the distributions of the $S_{MN}, ACV, AHP, R, I_{th}$ and $\tau$ properties between the normalized global datasets built in this study. For each property, the Range, Mean, coefficient of variation ($CoV = \frac{standard\ deviation}{Mean}$) and the ratio $\frac{Me}{Md} = \frac{Mean}{Median}$ of its distribution in a global dataset is compared to the other global datasets it appears in. For each property, the standard deviation $sd_G$ between global datasets of these four metrics is computed. A low $sd_G$ value reflects the low variability of the property distribution between global datasets.

**Table TSM1. $r^2$ values obtained for each experimental dataset $\{A; B\}$ when performing a linear regression analysis on $\{A; B\}$ directly ('Linear') and on the $ln(A) - ln(B)$ ('Power') and $ln(A) - B$ ('Exponential') transformations of $\{A; B\}$.** The $r^2$ values returned by the three types of regression cannot be directly compared to estimate the best model. However, the power fit returned $r^2 > 0.5$ for relatively more experimental datasets than the linear and exponential fits.

| | Studies | Linear | Power | Exponential |
|---|---|---|---|---|
| $\{ACV; S_{MN}\}$ | (Cullheim, 1978) | 0,42 | 0,45 | 0,43 |
| | (Kernell & Zwaagstra, 1981) | 0,62 | 0,63 | 0,6 |
| | (Burke et al., 1982) | 0,23 | 0,26 | 0,24 |
| $\{AHP; S_{MN}\}$ | (Zwaagstra, B. & Kernell, 1980) | 0,33 | 0,37 | 0,35 |
| $\{AHP; ACV\}$ | (Eccles et al., 1958) | 0,53 | 0,54 | 0,54 |
| | (Zwaagstra, B. & Kernell, 1980) | 0,42 | 0,44 | 0,42 |
| | (Gustafsson & Pinter, 1984a) | 0,71 | 0,68 | 0,7 |
| | (Foehring et al., 1987) | 0,15 | 0,14 | 0,14 |
| $\{R; S_{MN}\}$ | (Kernell & Zwaagstra, 1981) | 0,68 | 0,71 | 0,7 |
| | (Burke et al., 1982) | 0,41 | 0,5 | 0,46 |
| $\{R; ACV\}$ | (Kernell, 1966) | 0,70 | 0,65 | 0,64 |
| | (Burke, 1968) | 0,28 | 0,31 | 0,31 |
| | (Kernell & Zwaagstra, 1981) | 0,68 | 0,71 | 0,71 |
| | (Fleshman et al., 1981) | 0,3 | 0,21 | 0,2 |
| | (Gustafsson & Pinter, 1984a) | 0,45 | 0,44 | 0,42 |
| | (Sasaki, 1991) | 0,68 | 0,68 | 0,66 |
| $\{R; AHP\}$ | (Gustafsson, B., 1979) | 0,43 | 0,41 | 0,42 |
| | (Gustafsson & Pinter, 1984a) | 0,71 | 0,68 | 0,67 |
| | (Foehring et al., 1987) | 0,33 | 0,32 | 0,33 |
| | (Pinter & Vanden Noven, 1989) | 0,66 | 0,53 | 0,59 |
| | (Sasaki, 1991) | 0,57 | 0,49 | 0,52 |
| $\{I_{th}; R\}$ | (Kernell, 1966) | 0,52 | 0,52 | 0,68 |
| | (Fleshman et al., 1981) | 0,28 | 0,34 | 0,38 |
| | (Gustafsson & Pinter, 1984b) | 0,63 | 0,83 | 0,83 |
| | (Zengel et al., 1985) | 0,49 | 0,6 | 0,62 |
| | (Munson et al., 1986) | 0,34 | 0,42 | 0,47 |
| | (Foehring et al., 1987) | 0,34 | 0,47 | 0,47 |
| | (Krawitz et al., 2001) | 0,32 | 0,49 | 0,44 |
| $\{I_{th}; ACV\}$ | (Kernell & Zwaagstra, 1981) | 0,23 | 0,24 | 0,24 |
| | (Gustafsson & Pinter, 1984b) | 0,4 | 0,52 | 0,5 |
| $\{I_{th}; AHP\}$ | (Gustafsson & Pinter, 1984b) | 0,55 | 0,72 | 0,69 |
| $\{C; R\}$ | (Gustafsson & Pinter, 1984a) | 0,55 | 0,57 | 0,57 |
| $\{C; I_{th}\}$ | (Gustafsson & Pinter, 1984a) | 0,23 | 0,34 | 0,24 |
| $\{C; AHP\}$ | (Gustafsson & Pinter, 1984a) | 0,25 | 0,26 | 0,26 |
| $\{C; ACV\}$ | (Gustafsson & Pinter, 1984a) | 0,17 | 0,22 | 0,19 |
| $\{\tau; R\}$ | (Burke & Ten Bruggencate, 1971) | 0,04 | 0,07 | 0,03 |
| | (Gustafsson, B., 1979) | 0,6 | 0,63 | 0,61 |
| | (Gustafsson & Pinter, 1984a) | 0,63 | 0,69 | 0,59 |
| | (Zengel et al., 1985) | 0,39 | 0,33 | 0,36 |
| | (Pinter & Vanden Noven, 1989) | 0,65 | 0,69 | 0,62 |
| | (Sasaki, 1991) | 0,68 | 0,72 | 0,64 |
| $\{\tau; I_{th}\}$ | (Gustafsson & Pinter, 1984b) | 0,63 | 0,59 | 0,57 |
| $\{\tau; ACV\}$ | (Gustafsson & Pinter, 1984a) | 0,69 | 0,76 | 0,76 |
| $\{\tau; AHP\}$ | (Gustafsson & Pinter, 1984a) | 0,34 | 0,32 | 0,31 |

**Table TSM2. Mathematical empirical NORMALIZED relationships between the MN properties $D_{soma}, R, C, \tau, I_{th}$, $AHP$ and $ACV$.** Each column provides the relationships between one and the six other MN properties. All constants and properties are normalized up to a theoretical 100% maximum value. To scale the normalized relationships $A = k \cdot B^c$ for a specific mammalian species, the normalized intercept $k$ must be scaled with a pair of values for properties $A$ and $B$ obtained from the same motoneuron in that species.

| | $D_{soma}$ | $R$ | $C$ | $\tau$ | $I_{th}$ | $AHP$ | $ACV$ |
|---|---|---|---|---|---|---|---|
| $D_{soma}$ | | $R = \frac{9.6 \cdot 10^5}{D_{soma}^{2.4}}$ | $C = 1.2 \cdot D_{soma}$ | $\tau = \frac{2.6 \cdot 10^4}{D_{soma}^{1.5}}$ | $I_{th} = 9.0 \cdot 10^{-4} \cdot D_{soma}^{2.5}$ | $AHP = \frac{2.5 \cdot 10^4}{D_{soma}^{1.5}}$ | $ACV = 4.0 \cdot D_{soma}^{0.7}$ |
| $R$ | $D_{soma} = \frac{2.9 \cdot 10^2}{R^{0.4}}$ | | $C = \frac{2.7 \cdot 10^2}{R^{0.4}}$ | $\tau = 5.8 \cdot R^{0.6}$ | $I_{th} = \frac{1.5 \cdot 10^3}{R}$ | $AHP = 4.7 \cdot R^{0.6}$ | $ACV = \frac{2.0 \cdot 10^2}{R^{0.3}}$ |
| $C$ | $D_{soma} = 8.6 \cdot 10^{-1} \cdot C$ | $R = \frac{1.4 \cdot 10^6}{C^{2.5}}$ | | $\tau = \frac{3.3 \cdot 10^4}{C^{1.5}}$ | $I_{th} = 6.2 \cdot 10^{-4} \cdot C^{2.6}$ | $AHP = \frac{3.1 \cdot 10^4}{C^{1.6}}$ | $ACV = 3.6 \cdot C^{0.7}$ |
| $\tau$ | $D_{soma} = \frac{9.5 \cdot 10^2}{\tau^{0.7}}$ | $R = 5.6 \cdot 10^{-2} \cdot \tau^{1.6}$ | $C = \frac{8.5 \cdot 10^2}{\tau^{0.7}}$ | | $I_{th} = \frac{2.9 \cdot 10^4}{\tau^{1.7}}$ | $AHP = 7.8 \cdot 10^{-1} \cdot \tau$ | $ACV = \frac{4.6 \cdot 10^2}{\tau^{0.5}}$ |
| $I_{th}$ | $D_{soma} = 1.6 \cdot 10^1 \cdot I_{th}^{0.4}$ | $R = \frac{1.1 \cdot 10^3}{I_{th}}$ | $C = 1.7 \cdot 10^1 \cdot I_{th}^{0.4}$ | $\tau = \frac{4.2 \cdot 10^2}{I_{th}^{0.6}}$ | | $AHP = \frac{3.7 \cdot 10^2}{I_{th}^{0.6}}$ | $ACV = 2.7 \cdot 10^1 \cdot I_{th}^{0.3}$ |
| $AHP$ | $D_{soma} = \frac{8.1 \cdot 10^2}{AHP^{0.7}}$ | $R = 8.4 \cdot 10^{-2} \cdot AHP^{1.6}$ | $C = \frac{7.3 * 10^2}{AHP^{0.6}}$ | $\tau = 1.3 \cdot AHP$ | $I_{th} = \frac{1.9 * 10^4}{AHP^{1.7}}$ | | $ACV = \frac{4.1 \cdot 10^2}{AHP^{0.5}}$ |
| $ACV$ | $D_{soma} = 1.4 \cdot 10^{-1} \cdot ACV^{1.4}$ | $R = \frac{1.2 \cdot 10^8}{ACV^{3.5}}$ | $C = 1.7 \cdot 10^{-1} \cdot ACV^{1.4}$ | $\tau = \frac{5.0 \cdot 10^5}{ACV^{2.1}}$ | $I_{th} = 5.9 \cdot 10^{-6} \cdot ACV^{3.6}$ | $AHP = \frac{5.0 \cdot 10^5}{ACV^{2.2}}$ | |

**Table TSM3. Typical ranges of physiological values for $D_{soma}$ and $S_{neuron}$ in cat rat and mouse species.** $S_{MN}$ is found to vary over an average $q_S^E = 2.4$-fold range, which sets the amplitude of the theoretical ranges. Absolute {min; max} reports the minimum and maximum values retrieved in the reference studies for $D_{soma}$ and $S_{neuron}$, while average {min; max} is obtained as the average across reference studies of minimum and maximum values retrieved per study.

| | Property | Unit | Absolute {min;max} | Average {min; max} | $q_S^E$ | Reference studies | Theoretical range |
|---|---|---|---|---|---|---|---|
| Cat | $D_{soma}$ | $[\mu m]$ | $\{25.0; 90.0\}$ | $\{36.7; 75.5\}$ | 2.2 | (Kernell, 1966; Cullheim, 1978; Zwaagstra, B. & Kernell, 1980; Kernell & Zwaagstra, 1981; Ulfhake & Kellerth, 1981; Zwaagstra & Kernell, 1981; Burke et al., 1982; Donselaar et al., 1986; Destombes et al., 1992) | $[33; 79]$ |
| | $S_{neuron}$ | $[mm^2]$ | $\{0.08; 0.64\}$ | $\{0.17; 0.45\}$ | 2.7 | (Barrett & Crill, 1974; Ulfhake & Kellerth, 1981; Burke et al., 1982; Ulfhake, B. & Kellerth, 1984; Ulfhake, Brun & Cullheim, 1988; Moschovakis et al., 1991) | $[0.18; 0.44]$ |
| Rat | $D_{soma}$ | $[\mu m]$ | $\{17.5; 66.0\}$ | $\{23.9; 55.7\}$ | 2.4 | (Swett et al., 1986; von Steyern et al., 1999; Copray & Kernell, 2000; Ishihara et al., 2001; Deardorff et al., 2013; Mierzejewska-Krzyżowska et al., 2014) | |
| Mouse | $D_{soma}$ | $[\mu m]$ | $\{14.0; 35.0\}$ | $\{14.0; 35.0\}$ | 2.5 | (von Steyern et al., 1999) | |
| | $S_{neuron}$ | $[mm^2]$ | $\{0.01; 0.08\}$ | $\{0.01; 0.06\}$ | 4.4 | (Amendola & Durand, 2008; Brandenburg et al., 2020) | |

**Table TSM4: Typical ranges of physiological values for the MN properties $\boldsymbol{R}, \boldsymbol{R_m}, \boldsymbol{C}, \boldsymbol{\tau}, \boldsymbol{I_{th}}, \boldsymbol{AHP}$ and $\boldsymbol{ACV}$.** As described in the Methods section, $q_A^E$ is the average among reference studies of the ratios of minimum and maximum values; the properties experimentally vary over a $q_A^E$-fold range. This ratio compares with the theoretical ratio $q_A^T = (q_S^E)^{|c|}$ (with $c$ taken from Table 3 in manuscript), which sets the amplitude of the theoretical ranges. Absolute and average {min; max} are obtained as described in the main manuscript.

| MN property | Unit | Absolute exp {min;max} | Average exp {min; max} | $q_A^E$ | Reference studies | $q_A^T$ | $\frac{q_A^E}{q_A^T}$ | Theoretical range |
|---|---|---|---|---|---|---|---|---|
| $\boldsymbol{R}$ | $M\Omega$ | $\{0.2; 8.1\}$ | $\{0.4; 4.0\}$ | 10.9 | (Kernell, 1966; Burke, 1968; Burke & Ten Bruggencate, 1971; Barrett & Crill, 1974; Gustafsson, B., 1979; Kernell & Zwaagstra, 1981; Fleshman et al., 1981; Burke et al., 1982; Ulfhake, B. & Kellerth, 1984; Gustafsson & Pinter, 1984a; Zengel et al., 1985; Foehring et al., 1986; Munson et al., 1986; Foehring et al., 1987; Sasaki, 1991; Krawitz et al., 2001) | 8.4 | 1.3 | $[0.5; 4.0]$ |
| $\boldsymbol{C}$ | $nF$ | $\{2.2; 8.5\}$ | $\{2.2; 8.5\}$ | 3.9 | (Gustafsson & Pinter, 1984a; Gustafsson & Pinter, 1985) | 2.3 | 1.6 | $[3.2; 7.5]$ |
| $\boldsymbol{\tau}$ | $ms$ | $\{2.0; 14.2\}$ | $\{2.9; 10.2\}$ | 3.5 | (Burke & Ten Bruggencate, 1971; Barrett & Crill, 1974; Gustafsson, B., 1979; Ulfhake, B. & Kellerth, 1984; Gustafsson & Pinter, 1984a; Gustafsson & Pinter, 1985; Sasaki, 1991) | 3.7 | 0.9 | $[2.8; 10.3]$ |
| $\boldsymbol{I_{th}}$ | $nA$ | $\{1.7; 52.7\}$ | $\{2.3; 36.6\}$ | 16.3 | (Fleshman et al., 1981; Kernell & Monster, 1981; Ulfhake, B. & Kellerth, 1984; Gustafsson & Pinter, 1984b; Zengel et al., 1985; Foehring et al., 1986; Munson et al., 1986; Foehring et al., 1987; Krawitz et al., 2001) | 9.1 | 1.8 | $[3.9; 35.0]$ |
| $\boldsymbol{AHP}$ | $ms$ | $\{10.6; 266.6\}$ | $\{44.2; 158.7\}$ | 4.1 | (Eccles et al., 1957; Gustafsson, B., 1979; Dum & Kennedy, 1980; Zwaagstra, B. & Kernell, 1980; Ulfhake, B. & Kellerth, 1984; Gustafsson & Pinter, 1984a; Zengel et al., 1985; Foehring et al., 1987; Sasaki, 1991) | 3.8 | 1.1 | $[42.6; 160.2]$ |
| $\boldsymbol{ACV}$ | $m \cdot s^{-1}$ | $\{51.1; 127.2\}$ | $\{65.3; 114.8\}$ | 1.8 | (Eccles et al., 1957; McPhedran et al., 1965; Kernell, 1966; Appelberg & Emonet-Dénand, 1967; Burke, 1968; Barrett & Crill, 1974; Proske & Waite, 1974; Bagust, 1974; Stephens & Stuart, 1975; Cullheim, 1978; Dum & Kennedy, 1980; Zwaagstra, B. & Kernell, 1980; Kernell & Zwaagstra, 1981; Fleshman et al., 1981; Glenn & Dement, 1981; Burke et al., 1982; Gustafsson & Pinter, 1984a; Zengel et al., 1985; Foehring et al., 1986; Foehring et al., 1987) | 2.3 | 0.8 | $[63.5; 116.6]$ |

**Table TSM5. Validation of the cat relationships (Table 6 in manuscript) against rat and mouse data**. nME: normalized maximum error. nRMSE: normalized root-mean-square error. $r^2_{pred}$: coefficient of determination between experimental and predicted quantities. $r^2_{exp}$: coefficient of determination of the power trendline directly fitted to the experimental data.

| Animal | Dataset | Reference studies | $nME$ (%) | $nRMSE$ (%) | $r^2_{pred}$ | $r^2_{exp}$ |
|---|---|---|---|---|---|---|
| **Rat** | $\{I_{th}; R\}$ | (Gardiner, 1993; Bakels & Kernell, 1993; Lee & Heckman, 1998; Button et al., 2008; Turkin et al., 2010; Krutki et al., 2015) | 352 | 16 | 0.53 | 0.54 |
| **Mouse** | $\{I_{th}; R\}$ | (Delestrée et al., 2014; De Lourdes Martínez-Silva et al., 2018; Huh et al., 2021) | 385 | 16 | 0.46 | 0.46 |
| | $\{\tau; R\}$ | (Manuel et al., 2009) | 1219 | 168 | 0.35 | 0.40 |
| | $\{C; R\}$ | (Manuel et al., 2009) | 1915 | 98 | 0.38 | 0.38 |
